# Regenerating motor neurons prime muscle stem cells for myogenesis by enhancing protein synthesis and mitochondrial bioenergetics

**DOI:** 10.1101/2020.05.24.113456

**Authors:** Jeongmoon J. Choi, Eun Jung Shin, Woojin M. Han, Shannon E. Anderson, Mahir Mohiuddin, Nan Hee Lee, Thu Tran, Shadi Nakhai, Hyeonsoo Jeong, Anna Shcherbina, Gunjae Jeong, Dong Gun Oh, Laura D. Weinstock, Sitara B. Sankar, Molly E. Ogle, Lida Katsimpardi, Tata Nageswara Rao, Levi Wood, Carlos A. Aguilar, Amy J. Wagers, Young C. Jang

## Abstract

Throughout life, skeletal muscle, the arbiter of voluntary movements, is maintained by a population of skeletal muscle-dedicated stem cells, called muscle satellite cells (MuSCs). Similar to other adult stem cells, the function of MuSCs is tightly coordinated by the cellular and acellular components of their microenvironment, or the niche. While the processes that control the coupling of neurotransmission and muscle contraction have been well characterized, little is known on the reciprocal crosstalk between neural cells and MuSCs within the muscle microenvironment. Here, we report that mild peripheral nerve injury enhances MuSC myogenic function and muscle regeneration by synergistically augmenting MuSC mitochondrial bioenergetics and upregulating anabolic protein synthesis pathways. We also demonstrate that chronic disruption or degeneration of neuromuscular synapses, such as in muscular dystrophy and biological aging, abolishes MuSC and motor neuron interactions, causing significant deficits in muscle regeneration following injury. These results underscore the importance of neuromuscular junction and neural network as an essential niche of MuSCs. Determining the significance of MuSC-nerve interactions and their functional outcomes, as well as the possibility of modulating these connections, have important implications for our understanding of neuromuscular disease pathology and development of therapeutic interventions.

**Highlights:**

- Mild peripheral nerve injury increases muscle stem cell bioavailability of healthy muscle.
- Nerve perturbation stimulates myogenesis by enhancing protein synthesis and mitochondrial metabolism in young, healthy muscle.
- Synergistic crosstalk within neuromuscular niche boosts muscle regeneration in young, healthy muscle.
- Positive influences from the neural network on muscle stem cells are abolished in pathological denervation manifested in dystrophic and aging muscle.

## INTRODUCTION

Skeletal muscle possesses remarkable regenerative capacity due to the presence of resident stem cells, called muscle satellite cells (MuSC) (Almada and Wagers, 2016; Mauro, 1961). During normal homeostasis, MuSCs remain in quiescence underneath the basal lamina and adjacent to myofibers. However, when there are detrimental alterations in physiology or injuries to the muscle microenvironment, MuSCs express sets of transcriptional factors that promote asymmetric division in which committed progenies differentiate and fuse with existing myofibers or form *de novo* myofibers, while other populations of MuSC progeny self-renew to replenish the quiescent stem cell pool for future rounds of regeneration (Almada and Wagers, 2016). These orchestrated sequences of cellular events play a crucial role in maintaining muscle plasticity and remodeling of multinucleated syncytia in response to external stimuli (Almada and Wagers, 2016; Bassel-Duby and Olson, 2006). Similar to other adult stem cells, a growing amount of evidence suggests that the MuSC function is also tightly coordinated by cellular and acellular components surrounding its microenvironment, also known as the MuSC niche (Almada and Wagers, 2016; Wagers, 2012; Yin et al., 2013).

A critical conduit through which MuSCs alter their state is via interactions with axons from motor neurons, which form synapses with skeletal muscle fibers at the neuromuscular junction (NMJ) (Lichtman and Sanes, 2003; Sanes and Lichtman, 1999). The neurotransmission from motor neurons not only controls muscle force generation through the excitation-contraction coupling but also regulates gene expression patterns of adult myofibers (Sanes and Lichtman, 1999). Perturbations of NMJ have been linked to several neuromuscular pathologies, including aging and muscular dystrophies. Affected muscles elicit muscle wasting and weakness (Jang and Van Remmen, 2011; Shi et al., 2014; Valdez et al., 2012), and genetic ablation of MuSCs have been shown to impair the regeneration of NMJ after sciatic nerve transection (Liu et al., 2017; Liu et al., 2015). Conversely, MuSCs in denervated muscle fail to terminally differentiate and form smaller myotubes (Borisov et al., 2005; Dedkov et al., 2003). Anatomically, a specific subset of MuSCs is located in close proximity to the NMJ, suggesting potential interactions between MuSCs and the NMJ (Liu et al., 2017; Liu et al., 2015; Relaix and Zammit, 2012). Moreover, small subpopulations of myonuclei are known to reside underneath NMJs (Ruegg, 2005; Zhang et al., 2007), are transcriptionally distinct from other myonuclei within the muscle fiber, and determine the size and synaptic activity of motor endplates (Ruegg, 2005; Zhang et al., 2007). Given that the primary source of these myonuclei is MuSCs, and both the NMJ and MuSCs are modified in aging and disease, these observations support the notion of a synergistic interaction between MuSCs and the NMJ. Yet, the mechanism through which neural activity and the NMJ influences MuSC actions has not been defined.

Here, we report that mild peripheral nerve injury boosts muscle satellite cell myogenic function and muscle regeneration by synergistically enhancing MuSC mitochondrial bioenergetics and upregulating anabolic protein synthesis pathways. We also demonstrate that chronic disruption or degeneration of neuromuscular synapses, such as in muscular dystrophy and biological aging, abolishes muscle stem cell and motor neuron interactions, causing significant deficits in muscle regeneration following injury. These results highlight the importance of reciprocal interactions between the neural network and MuSCs during muscle regeneration and remodeling.

## RESULTS

### Peripheral nerve injury increases MuSC bioavailability

To examine how MuSC fate and myogenic progression are modified in response to perturbation of motor neurons, we employed sciatic nerve crush (SNC) injury (axonotmesis), in which motor neurons undergo Wallerian degeneration while epineurium is left intact (Bauder and Ferguson, 2012; Ma et al., 2011). Similar to previously reported studies, pre-synaptic motor axons were retracted and denervated within 7 days and start to re-occupy synapses by day 14 following injury (Bauder and Ferguson, 2012; Kang and Lichtman, 2013; Kang et al., 2014) (**Fig. 1A**). In stark contrast to cardiotoxin injury, in which both presynaptic motor axon and post-synaptic acetylcholine receptors (AChR) are altered in SNC (**Fig. S1A-1C**), the AChR remained intact, and only the pre-synaptic sides were retracted (**Fig. 1A**, *bottom*). Next, to assess changes in MuSCs following SNC, we quantified the total number of quiescent Pax7-expressing MuSCs on single myofibers isolated from sham and denervated tibialis anterior (TA) muscle 7 days after injury. Denervated muscle showed a significant increase in Pax7^+^ MuSCs compared to contralateral controls (administered sham-surgery) (**Fig. 1B, C**). To confirm the elevation of MuSC frequency in denervated muscle, we analyzed FACS purified MuSCs from MuSC reporter mice that express *ZsGreen* fluorescent protein under the control of *Pax7* regulatory element, as well as the double sorted population from hindlimb of young wildtype mice using well-established MuSC surface markers (CD45^−^/CD11b^−^/CD31^−^/Ter119^−^/Sca1^−^/β1-integrin^+^/CXCR4^+^)(Cerletti et al., 2008; Jang et al., 2011; Sherwood et al., 2004). In both cases, when normalized to the weight of the muscle used, we found an approximately 50% increase in MuSC yield from denervated myofibers. We next compared the ultra-structure of freshly isolated MuSCs from control and 7-day denervated hindlimb muscles and observed differences in cellular morphology whereby denervated MuSCs display prevalence of euchromatin and concomitant reductions of heterochromatin (**Fig. 1F** *bottom*). Furthermore, z-stacked confocal images and 3D tomographic images showed denervated MuSCs significantly increased average cell volume (**Fig. 1F, G**). These results suggest denervated MuSCs may have transitioned to a G_alert_ state, which is characterized by an increase in cell volume with high mitochondrial content and accelerated activation and proliferation kinetics (Rodgers et al., 2014; Rodgers et al., 2017).

**Figure 1.**
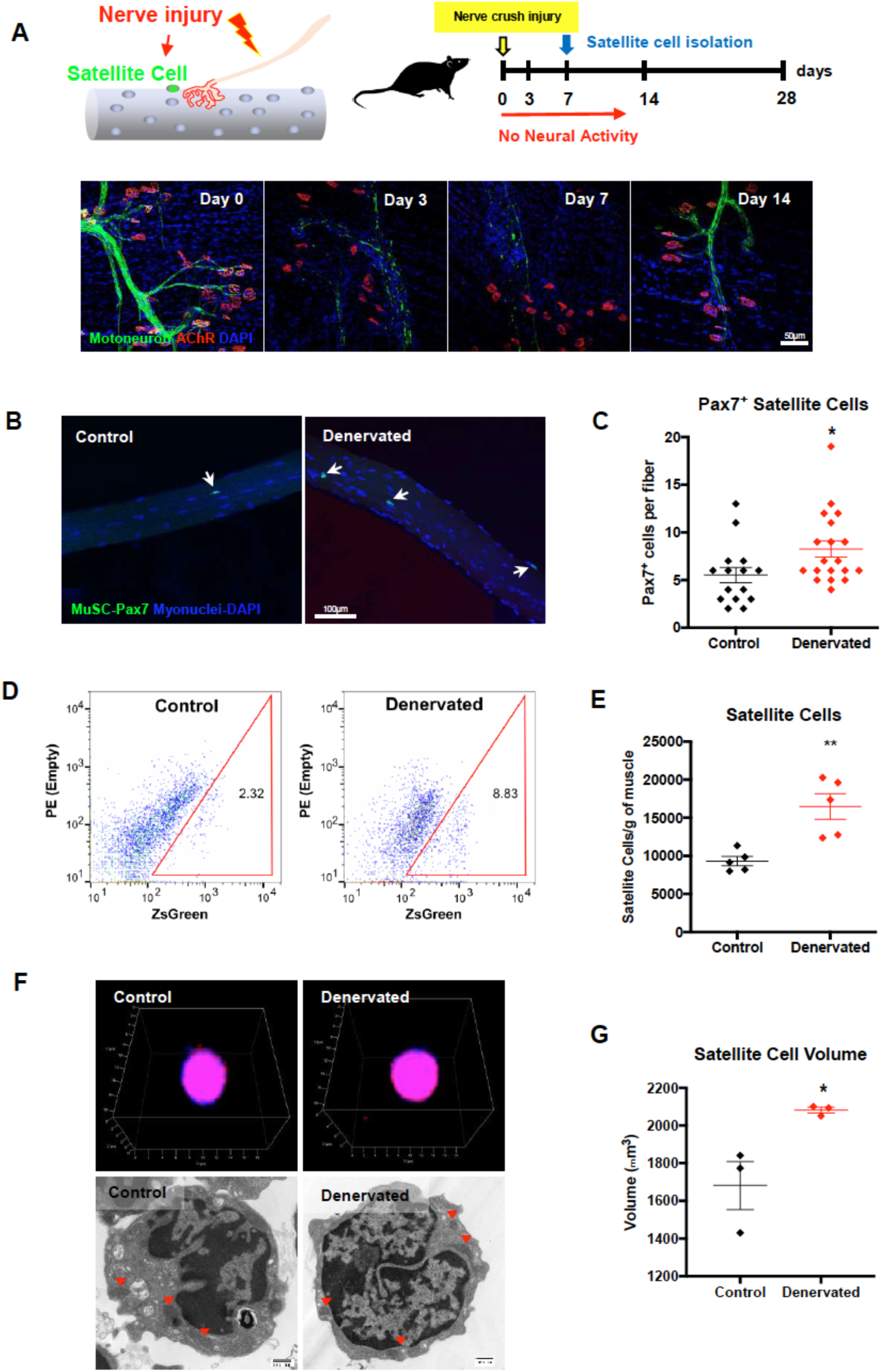
Peripheral nerve injury increases satellite cell content. **(A)** Schematics of experimental design and nerve regeneration timeline following pinched-sciatic nerve injury. **(B)** Representative images of denervated single myofibers showing increased Pax7+ satellite cells. **(C)** Quantification of satellite cells content comparing sham control and 7 days following sciatic nerve pinch injury. \**p*<0.05, Mean ± SEM **(D)** Representative FACS plot of satellite cell frequency from control and denervated *Pax7-ZsGreen*, muscle satellite cell reporter mice. **(E)** Quantification of FACS purified satellite cell yield normalized to muscle mass. \*\**p*<0.01, Mean ± SEM **(F)** Representative z-stack confocal images of single satellite cell (*top*) and transmission electron micrograph (TEM) of satellite cells (*bottom*). Red arrowheads indicate mitochondria **(E)** Quantification of freshly isolated satellite cell volume 7 days following denervation. \**p*<0.05, Mean ± SEM (Scale bars; (A) 50 µm, (B) 100 µm, (F) 500 nm)

### MuSCs are primed for myogenic differentiation in response to nerve crush injury

To determine the transcriptional changes associated with acute nerve injury, we isolated MuSCs using fluorescence-activated cell sorting (FACS) from 7-day denervated muscle and contralateral control and analyzed MuSC transcriptome using RNA-sequencing (**Fig. 2**). Unsupervised clustering analysis showed prominent differences in the transcriptome between control and 7-day denervated quiescent MuSCs, with over 1,000 differently expressed genes (top one hundred genes are shown in **Fig. 2A**). Gene set enrichment analysis (GSEA) and pathway analysis of differentially expressed genes identified numerous pathways associated with denervation including striated muscle contraction, muscle development, cell cycle regulation, p38 MAPK, and metabolism (**Fig. 2B, C**). Consistent with G_alert_ state, analysis of the highly expressed and differentially expressed transcripts suggest that SNC injury primes satellite cells towards myogenic progression and contractile muscle formation (**Fig. S2A-2C**).

**Figure 2.**
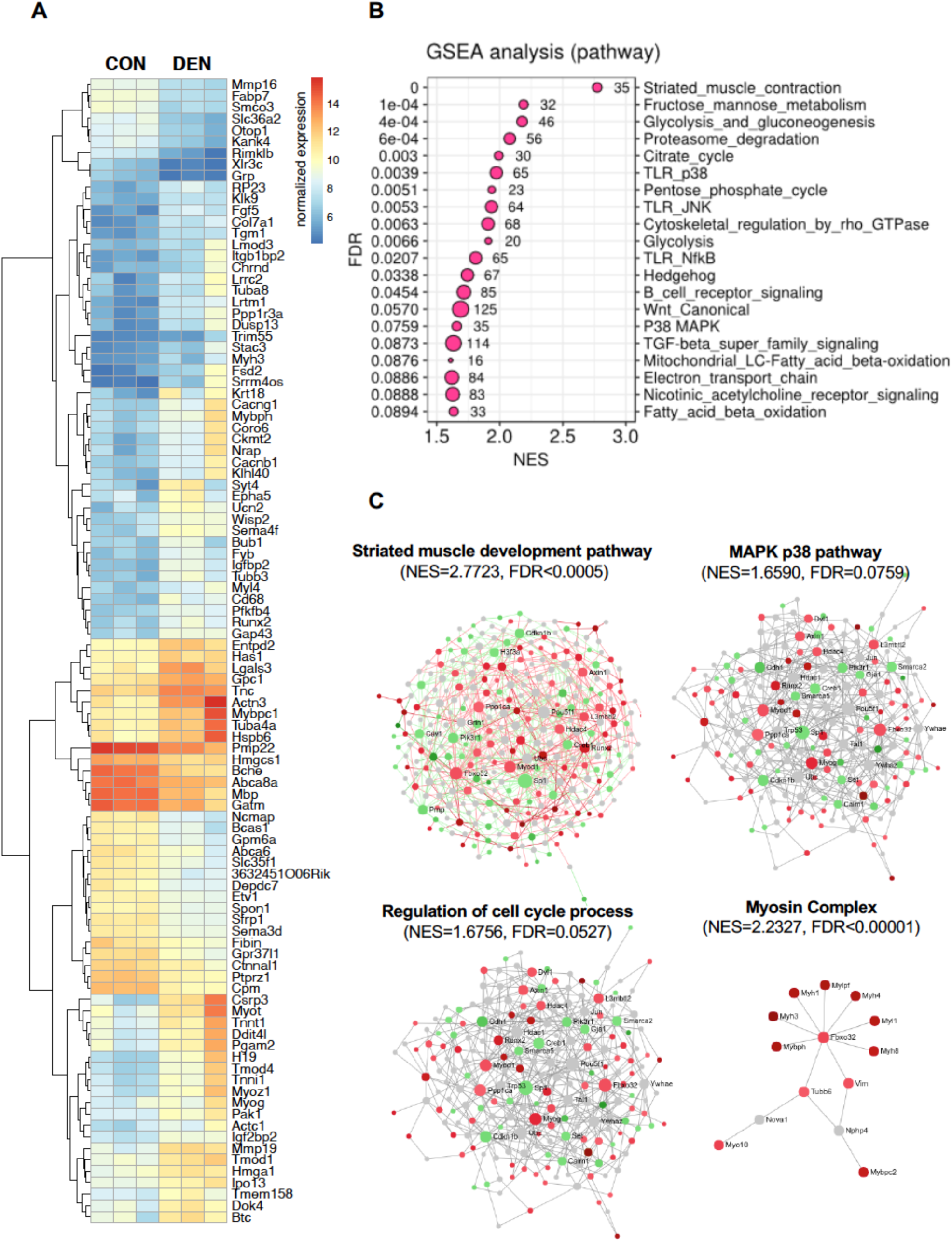
Transcriptomic analyses show enhanced myogenic and metabolic signatures in denervated muscle satellite cells. **(A)**Hierarchical clustering and heatmap representation (TPM, transcripts per million) of the top hundred most differentially expressed enriched genes (*p*<0.05, n=3). **(B)** Gene set enrichment analysis (GSEA) of top twenty pathways enriched in muscle satellite cells 7 days following denervation. **(C)** Pathways and gene interaction network analyses for striated muscle development, p38 MAPK, cell cycle, and myosin complex gene sets by gene set enrichment analysis comparison of control to denervated muscle satellite cells. Data were derived from three RNA-seq biological replicates. NES, normalized enrichment score; FDR, false discovery rate.

### Nerve injury stimulates myogenesis and protein synthesis *ex vivo*

To further examine changes in MuSC dynamics following nerve injury, MuSCs were FACS-purified 7 days after SNC injury and evaluated for myogenic changes. First, we assessed whether SNC injury altered MuSC surface marker used for flow cytometry (CD45^−^/CD11b^−^/CD31^−^/Ter119^−^/Sca1^−^/β1-integrin^+^/CXCR4^+^) by exploiting dimensionality reduction technique, SPADE (spanning-tree progression analysis of density-normalized events) (Qiu, 2017; Qiu et al., 2011). Representative frequency distribution of all live cells within the SPADE tree indicate differing cell populations exist in control versus SNC injury (**Fig. S3A**). SPADE analysis of the MuSC population produced 4 clusters of nodes that differentially expressed β1-integrin and CXCR4 (**Fig. S3B**). MuSCs from denervated muscle were significantly more represented in clusters of β1-integrin^low^/CXCR4^low^ populations versus control (**Fig. S3C**) (Rocheteau et al., 2012; Rozo et al., 2016). Accordingly, RNA-seq data showed that denervated MuSCs maintained similar levels in early MuSC activation markers (i.e., *Pax7, CD34, Sdc4, Myf5*) and cell cycle regulators, myogenic differentiation and myofibril assembly markers, such as *Myogenin, Myf6, Myh1, Acta1, Tnnt1, and Myoglobin*, were significantly upregulated compared to contralateral controls (**Fig. 3A**). These results further support the notion that denervated MuSCs are transcriptionally primed to engage in myogenesis, similar to the transition from G_0_ to G_alert_ state (Rodgers et al., 2014; Zhang et al., 2015). Next, to corroborate an enhanced myogenic activity of denervated MuSCs, we assessed proliferation and differential potential. As expected, when cells were cultured *ex vivo* for 3 days, MuSCs from nerve injury showed an approximately 2-fold increase in expansion, as measured by total myoblast number (**Fig. 3C**). Consistent with a higher frequency of myoblasts, denervated MuSCs exhibited increased proliferation as measured by incorporation of 5-ethynyl-2’-deoxyuridine (EdU) following 24 hours pulse-chase (**Fig. 3D**). Furthermore, when single MuSC were seeded and grown in 96-well plates, nerve-injured MuSCs formed colonies with up to 50% increased efficiency as compared to control cells, indicating that denervated muscle contains a substantially higher pool of functional regenerative cells (**Fig. 3E**). To further validate the improved myogenic activity of nerve-injured MuSCs, we next measured the differentiation of these cells by reducing serum concentration. When an equal number of myoblasts were seeded in a 2% serum condition, denervated MuSCs formed multinucleated, myosin heavy chain (MHC) expressing myotubes at a much faster rate compared to control cells (**Fig. 3B**, *bottom*). Likewise, the rate of fusion was significantly greater in denervated myoblasts. To corroborate the fusion capacity, we purified control MuSCs from cyan fluorescence protein (CFP) and yellow fluorescence protein (YFP) transgenic mice, and denervated MuSCs from TdTomato transgenic mice. An equal number of myoblasts in 3 different colors were randomly seeded and allowed to differentiate for 5 days, then the fluorescence of myotubes was quantified. When CFP^+^(control) and YFP^+^ (control) myoblasts were differentiated, cyan, green, and yellow myotubes were evenly distributed, 33%, 23%, and 44%, respectively. In stark contrast, when CFP^+^ (control) and TdTomato^+^ (denervated) myoblasts were seeded together, 61% were TdTomato^+^ and only 14% were magenta^+^ (CFP + TdTomato), suggesting denervated myoblasts were fusing at a higher rate compared to controls (**Fig. 3F**).

**Figure 3.**
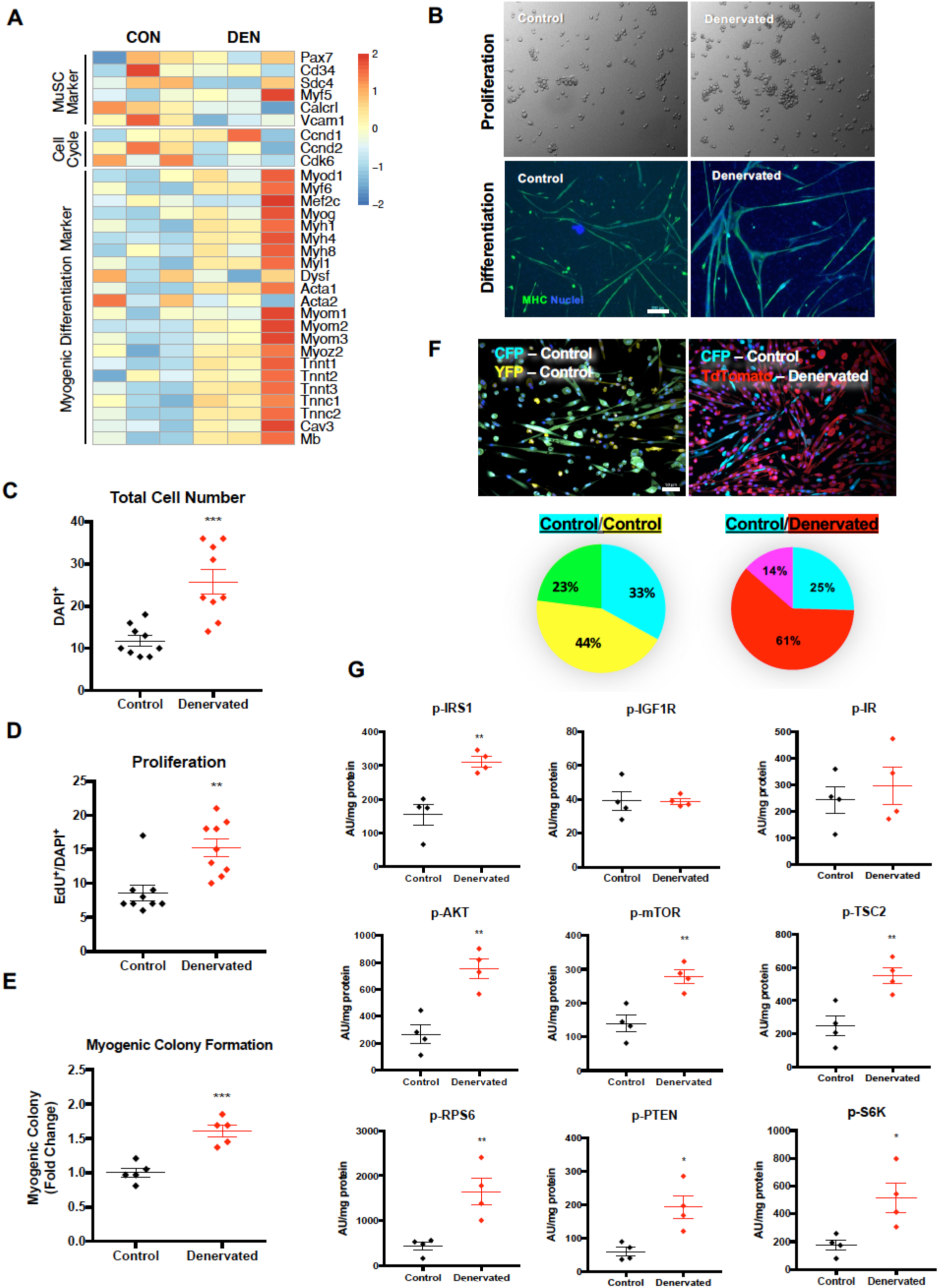
Nerve injury stimulates satellite cell proliferation, differentiation, and protein synthesis *ex vivo*. **(A)**Heatmap of key differentially expressed genes associated with satellite cell identity, cell-cycle, and myogenic differentiation. **(B)** Representative images of FACS purified MuSCs proliferation (*top*) and differentiation (*bottom*) in culture. Scale bar 200 mm. **(C)** Quantification of total satellite cells per cluster 48 hours after seeding. \*\*\**p*<0.001, Mean ± SEM, n=3 biological replicates. **(D)** Quantification of proliferating satellite cells 48 hours after seeding as measured by EdU^+^ and DAPI^+^ cells. **(E)** Single cell clonal expansion assays of sorted satellite cells from control or denervated mice. \*\*\**p*<0.001, Mean ± SEM. **(F)** Representative images of myogenic fusion assay from control (cyan fluorescent protein and yellow fluorescence protein) or denervated (TdTomato) mice (*top*). Scale bar 50 mm. Quantification of fusion rate in percentage (*bottom*). Data from n=3 biological replicates. **(G)** Luminex multiplex assay for protein synthesis pathway (phosphorylated IRS1, IGF1R, IR, AKT, mTOR, TSC2, RPS6, PTEN, and S6K) following 5 days in culture. \**p*<0.05, \*\**p*<0.01, Mean ± SEM, n=4 (Scale bars; (B) 200 µm, (F) 50 µm)

Since activated MuSCs are characterized by metabolic changes associated with increased biosynthesis of macromolecules (Almada and Wagers, 2016; Jang et al., 2011; Rodgers et al., 2014), and these changes are coordinated through the mammalian target of rapamycin (mTOR) signaling pathway (Bassel-Duby and Olson, 2006; Hornberger et al., 2006), we used a Luminex multiplex assay for anabolic protein synthesis. In concert with upregulation in genes associated with muscle formation and increased myogenesis, the phosphorylated (activated) insulin receptor substrate 1 (IRS-1), AKT, PTEN, mTOR, TSC2, S6K, RPS-6 protein contents were significantly elevated in nerve-injured myotubes compared to sham control myotubes (**Fig. 3G**). Taken together, these data further validate that SNC injury enriches MuSC population toward myogenesis progression.

### Mitochondrial metabolism in MuSCs is elevated in response to nerve damage

To further examine whether enhanced myogenesis and protein synthesis signaling seen in denervated MuSCs are coupled with elevated bioenergetics (Jang et al., 2011; Rodgers et al., 2014; Ryall et al., 2015; Tang and Rando, 2014), we assessed a series of metabolic parameters. First, in our GSEA, we found that citric cycle, glycolysis, mitochondrial fatty acid oxidation, electron transport chain, and pentose phosphate cycle gene sets were among highly upregulated pathways in nerve-injured MuSCs (**Fig. 2B**). In concert, we observed an upregulation of key mitochondrial citric acid cycle genes, *Aco2* and *Mdh2*, as well as nuclear-encoded mitochondrial ETC subunit genes, *Cyc1, Uqcrc1, Atp5g1*, and *Atp6c1b2* in MuSCs isolated from 7-day denervated muscle (**Fig. 4A**). Next, to evaluate whether mitochondrial morphology was different in denervated MuSCs, we utilized mitochondrial reporter mice, in which the fluorescent protein Dendra2 is targeted to mitochondrial matrix (Pham et al., 2012). While control MuSC mitochondria displayed elongated shape and concentrated near the nucleus, the majority of denervated MuSC mitochondria exhibit a punctate, fragmented shape with higher fission to readily support energetics for cell division (**Fig. 4B**, *top* **and Fig. S5A**). Moreover, in line with increased myogenic differentiation, denervated MuSCs show a significant expansion and higher mitochondrial content during fusion (**Fig. 4B**, *bottom*). Next, to assess whether changes in mitochondrial dynamics lead to alterations in bioenergetic function, we compared cellular respiration of control and 7-day denervated MuSCs in culture. Following 3 days of proliferation in growth media, cells were analyzed for mitochondrial oxygen consumption rate (OCR). As anticipated, denervated myoblasts showed higher basal OCR compare to control even when normalized to total protein concentration to account for the difference in cell number (**Fig. 4C**). In addition, when uncoupling agent, carbonyl cyanide m-chlorophenylhydrazone (CCCP) was added to measure maximal respiration, approximately a 2-fold increase was observed in nerve-injured myoblasts compared to controls (**Fig. 4C**), indicating bioenergetic function is synchronously enhanced with biosynthesis during myogenesis in denervated MuSCs. Despite changes in mitochondrial oxidative phosphorylation, we did not detect a significant difference in mitochondrial membrane potential, as measured by Tetramethyl rhodamine (TMRE) (**Fig. S5B**).

**Figure 4.**
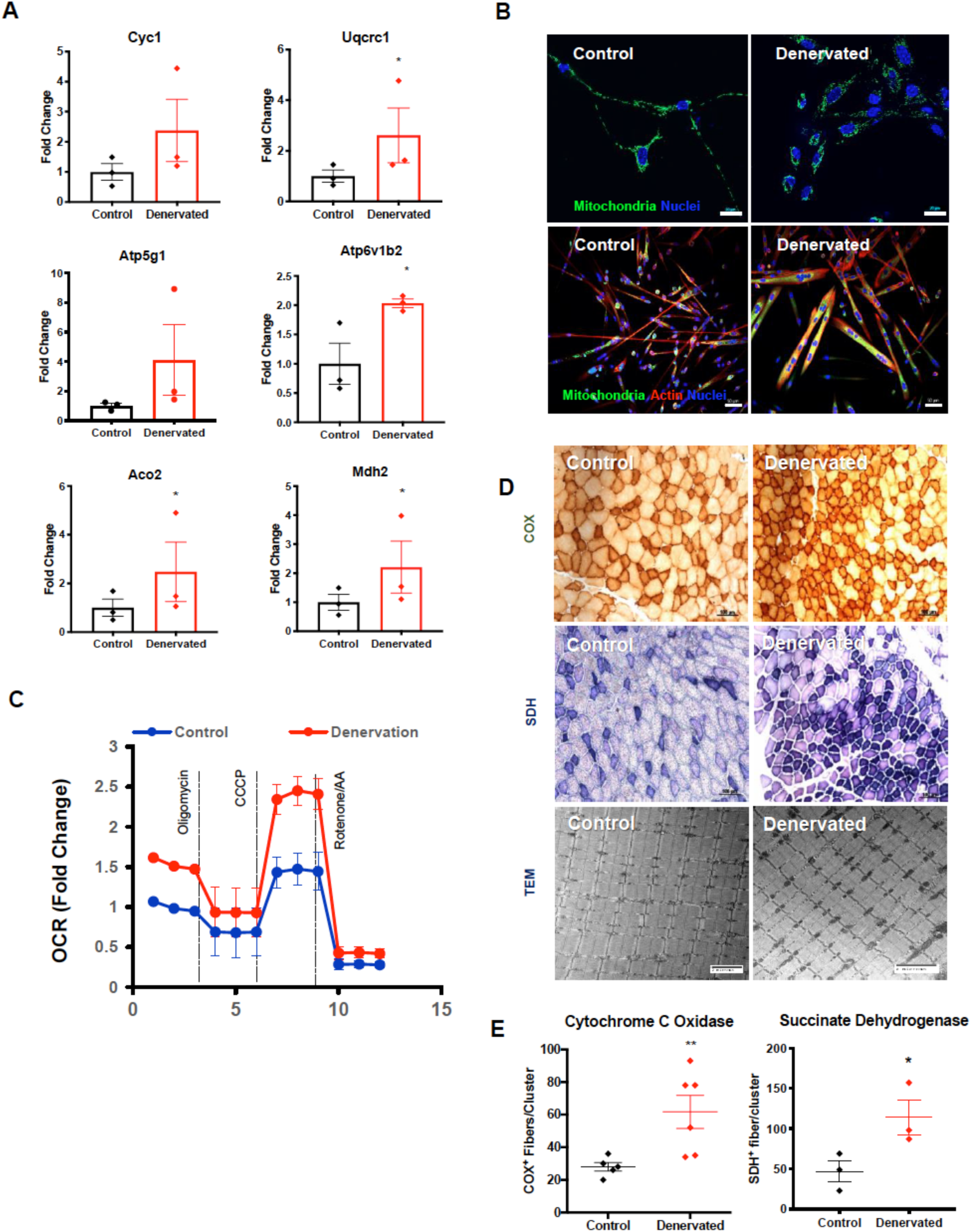
Nerve injury enhances satellite cell mitochondrial bioenergetics. **(A)**Mitochondrial electron transport chain (ETC) and citric acid cycle gene expression measured by qRT-PCR. **(B)** Representative images of MuSC mitochondrial morphology during proliferation (*top*) and differentiation (*bottom*). **(C)** Oxygen consumption rate (OCR) measurement from control and denervated muscle satellite 5 days after seeding, n=6. **(D)** Representative images of cytochrome c oxidase (COX; *top*), succinate dehydrogenase (SDH; *middle*), and TEM of interfibrillar mitochondria (*bottom*) from control and denervated tibialis anterior muscle 7 day following nerve injury. **(E)** Quantification of COX and SDH shown in **(D)**. \*\**p*<0.01, Mean ± SEM (Scale bars; (B) top 20 µm, bottom 50 µm, (D) top and middle 100 µm, bottom 2 µm)

During muscle regeneration, differentiated myotubes fuse to existing muscle or form *de novo* myofibers. Hence, we further asked whether enrichment in mitochondrial bioenergetics *ex vivo* also translated *in vivo* following denervation. Similar to findings from previous studies (Bonaldo and Sandri, 2013; Carlson, 2014; Langer et al., 2020), SNC injury induced progressive muscle atrophy in the lower hindlimb (**Fig. S6A-E**). To evaluate oxidative capacity of myofibers following SNC injury, succinate dehydrogenase (SDH) and cytochrome c oxidase (COX) activities were measured histologically. Similar to what we found in primary myoblasts, denervated muscle showed approximately 2-fold increase in the number of SDH^+^ and COX^+^ fibers 7 days after denervation (**Fig. 4D-E**). Consistent with these observations, Western blot analyses of the mitochondrial electron transport chain also revealed that SDH B subunit protein content was significantly higher in the denervated muscle homogenates (**Fig. S5C)**. Collectively, these data provide strong evidence that regenerative responses from nerve injury, prime MuSCs for myogenesis, and this enhancement is coupled with increased biosynthesis of myofiber proteins and mitochondrial bioenergetics.

### Nerve perturbation promotes a positive signaling microenvironment that augments muscle regeneration

We posited that the positive enhancements in MuSCs emanating from nerve injury would result in concomitant enhancements in muscle regeneration. Thus, we tested injury response from a composite injury, in which SNC was combined with muscle cryo-injury to tibialis anterior (TA) and compared it with muscle cryo-injury only in the contralateral TA, 7, 14, and 28 days after injury (**Fig. 5A**). In all of three timepoints analyzed, the regenerative potential, as measured by the number centrally nucleated myofibers and the expression of embryonic myosin heavy chain (eMHC) positive fibers, were significantly greater in nerve and muscle composite injury compared to muscle injury only group (**Fig. 5B, C**). Next, to corroborate enhanced muscle regeneration in the nerve-injured muscle, we transplanted an equal number GFP^+^ donor MuSCs into nerve plus muscle injury and muscle injury only recipients. 28 days following transplantation, TA muscles were harvested, and analyzed for engraftment efficiency (**Fig. 5D**). Consistent with enhanced regeneration, host muscle with denervation displayed a significantly higher number of GFP^+^ myofibers suggesting nerve injury synergistically augmented myogenesis of exogenous MuSCs (**Fig. 5E, F**). Summing these results demonstrates that changes in the regenerative environment from nerve injury promote MuSC regenerative dynamics.

**Figure 5.**
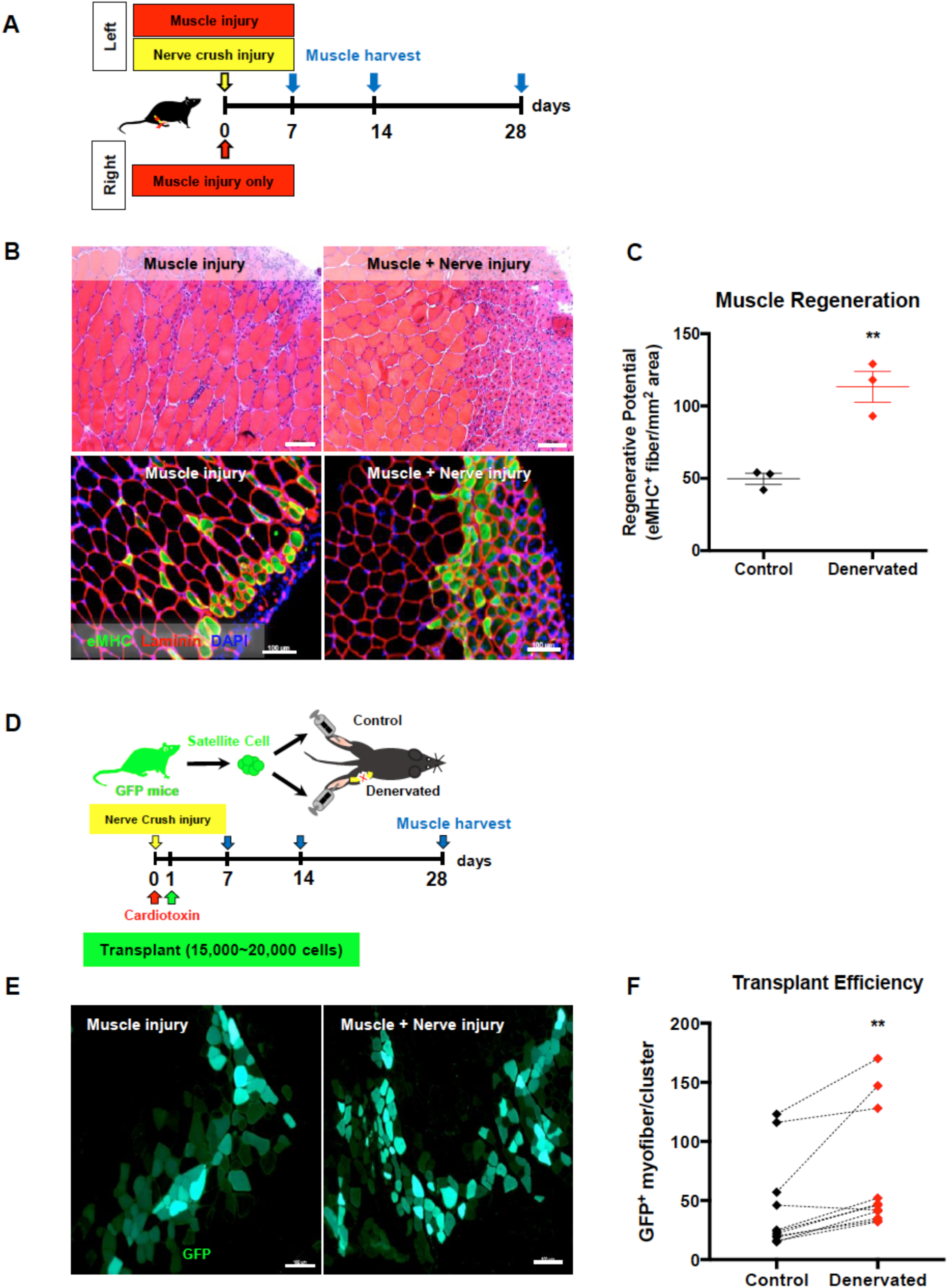
Nerve regeneration synergistically enhances skeletal muscle regeneration *in vivo*. **(A)** Schematic diagram of experimental design measuring muscle regeneration *in vivo*. **(B)** H&E (*top*) and immunofluorescence staining (embryonic myosin heavy chain, laminin, DAPI (*bottom*)) cross section from TA muscle of 7-day cardiotoxin injury (right leg) and cardiotoxin + sciatic nerve pinch injury (left leg). **(C)** Quantification of muscle regeneration assessed by the eMHC^+^ myofibers, \*\**p*<0.01, Mean ± SEM. **(D)** Schematic diagram of experimental design assessing transplantation efficiency. **(E)** Representative images of transplanted GFP^+^ fibers from muscle injury only and muscle and nerve injury. **(F)** Quantification of transplantation efficiency measured by total number of GFP^+^ fibers. \*\**p*<0.01, Mean ± SEM, n=10 (Scale bars; (B), (E) 100 µm)

### Positive interactions between motor neurons and MuSCs are lost in Duchenne muscular dystrophy

Duchenne muscular dystrophy (DMD) is a debilitating muscle disorder characterized by a loss of function mutation in dystrophin in the Sarcoglycan complex and concurrent breakdown of acetylcholine receptors (AChR) of the neuromuscular junction (NMJ) (Pratt et al., 2015). To explore whether synergistic interactions between motor neurons and MuSCs were intact in dystrophic muscle, we induced SNC injury to *mdx*, a mouse model of DMD, and analyzed MuSC function 7 days after injury. Consistent with reported studies(Pratt et al., 2015), motor endplate fragmentation was observed in all of the *mdx* hindlimb muscles analyzed, independent of injury, suggesting denervation was prevalent in dystrophic muscle (**Fig. 6A, B, S8**). Next, we tested if MuSC function and myogenic activation were augmented following SCN injury in *mdx* muscle. Unlike WT, hindlimb muscles from *mdx* mice failed to show an increase in MuSCs following 7-day denervation (**Fig. 6C)**. Similarly, RNA seq data revealed that the upregulation of key myogenesis genes following nerve injury was completely absent in dystrophic muscle (**Fig. 6D)**. Clustering analysis also exhibited noticeable differences in the transcriptome between WT and *mdx* MuSC in response to a 7-day SNC injury (**Fig. S9)**. Moreover, when sham and denervated MuSCs from *mdx* muscle were seeded *ex vivo*, there were no differences in proliferation or differentiation, suggesting that pro-regenerative signals emanating from neural microenvironment were ablated in dystrophic muscle (**Fig. 6E)**. We reasoned the lack of synergy after nerve injury was derived from intrinsic defects in mitochondrial functions of MuSCs, which have been previously reported (Percival et al., 2013; Schuh et al., 2012; Timpani et al., 2015). The basal mitochondrial respiration in WT and *mdx* MuSCs were compared and showed a significantly lower level of OCR for MuSCs from *mdx* muscle, even when normalized to total protein concentration, indicating an impaired metabolic capacity of *mdx* MuSCs (**Fig. 6F)**. Next, to further assess whether MuSC bioenergetics are enhanced following mild SNC injury, we applied the same 7-day nerve injury and measured OCR in FACS-purified MuSCs from sham-operated and denervated muscles. Unlike WT MuSCs in which nerve injury prompted elevated mitochondrial respiration, nerve injury in *mdx* MuSCs failed to promote mitochondrial OCR both in basal and maximal respiration (**Fig. 6G)**. Furthermore, metabolic pathway analysis from RNA seq data showed gene sets that were significantly upregulated following denervation in WT were not evident in *mdx* MuSCs. Importantly, genes in the mitochondrial electron transport chain (ETC) were noticeably upregulated in WT, but in stark contrast, they were downregulated in *mdx* MuSCs (**Fig. 6H)**. Lastly, to test synergistic improvement in the regenerative capacity of *mdx* mice, we induced concurrent sciatic nerve crush and cryo-injury in one leg, and only cryo-injury in the contralateral leg. 7 days following injury, immunofluorescence analyses were performed to measure muscle regeneration by quantifying eMHC^+^ and centrally nucleated fibers. In accordance with the lack of myogenic and bioenergetic improvements in *mdx* MuSCs, there were no differences in the regenerative capacity of *mdx* muscle, suggesting pro-regenerative nerve-muscle interaction is deficient in *mdx* muscle (**Fig. 6I)**. These data all support the notion that the absence of dystrophin and associated chronic denervation in DMD abolish the regenerative microenvironment from a healthy peripheral neural network.

**Figure 6.**
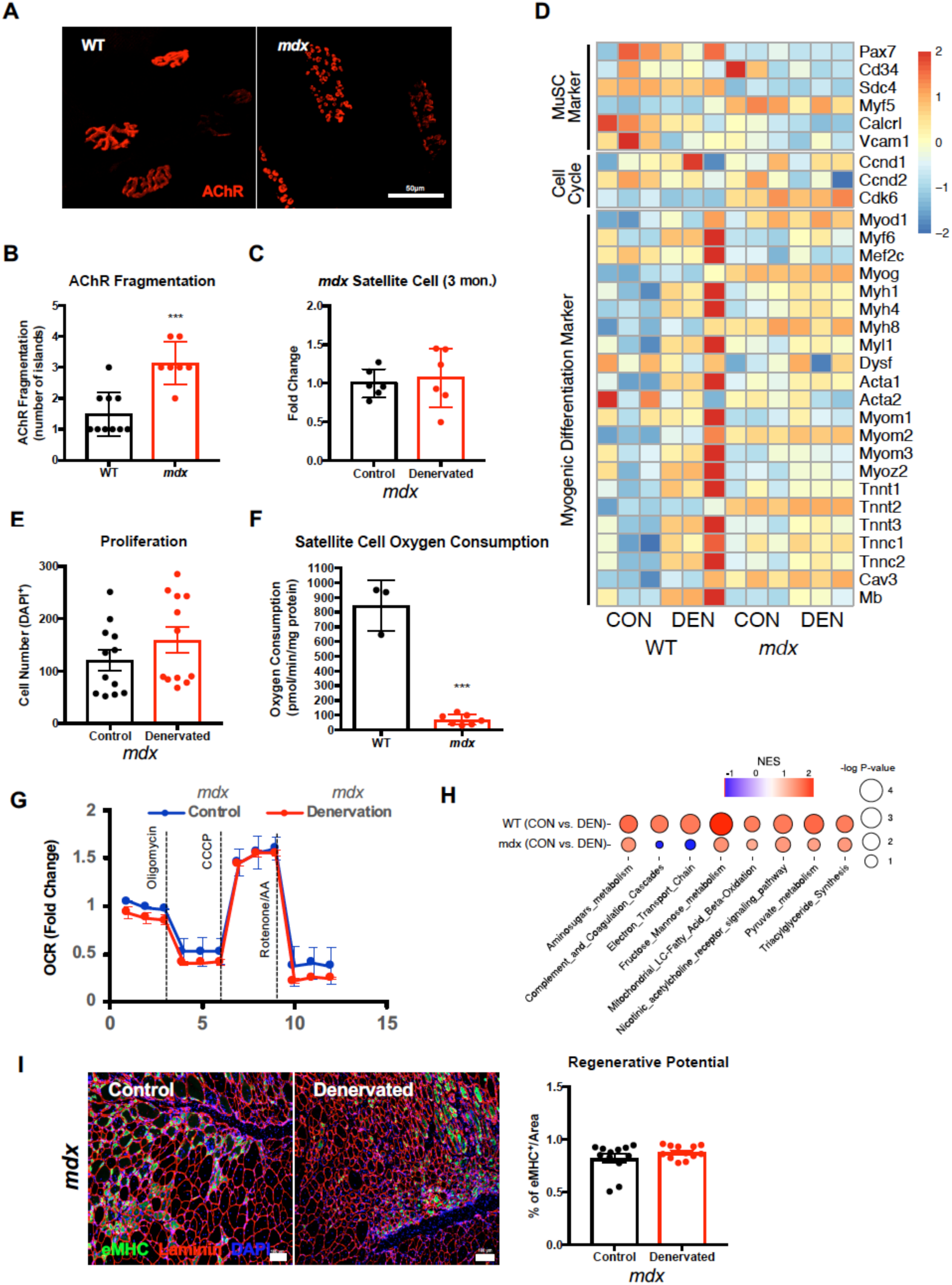
Satellite cell and motor neuron interactions are abolished in dystrophic muscle. **(A)** Representative images of wild-type and *mdx* neuromuscular junction, post-synaptic acetylcholine receptor stained with a-bungarotoxin. **(B)** Quantification of fragmented AChR. \*\*\**p*<0.001, Mean ± SEM. **(C)** Muscle satellite cell frequency from wildtype and mdx 7 days after denervation. Data expressed as Mean ± SD. **(D)** Heatmap of key differentially expressed genes associated with satellite cell identity, cell-cycle, and myogenic differentiation. **(E)** Quantification of total myoblasts 48 hours after seeding. **(F)** Satellite cell oxygen consumption rate from WT and *mdx*. \*\*\**p*<0.001, Mean ± SEM. **(G)** Oxygen consumption rate from muscle stem cell isolated from *mdx* and *mdx* 7-day denervation. **(H)** Dot plot showing gene set enrichment analysis (GSEA) of metabolic pathways differentially regulated in muscle satellite cells (WT *vs*. WT denervation and *mdx vs. mdx* denervation). **(I)** Representative immunofluorescence staining of embryonic myosin heavy chain, laminin, DAPI cross section from TA muscle of mdx 7-day cardiotoxin injury (right leg) and *mdx* cardiotoxin + sciatic nerve pinch injury (left leg). Quantification of regenerative potential measured by eMHC^+^ fibers shown in right. (Scale bars; (A) 50 µm, (I) 100 µm)

### Pro-regenerative interactions between motor neurons and MuSCs are absent in aged muscle

It has been well documented that skeletal muscle aging is associated with a progressive degeneration of NMJs and loss of motor units (Jang et al., 2011; Jang and Van Remmen, 2011; Liu et al., 2017). To extend our previous findings, we examined whether positive interactions from neuromuscular niche are altered as a function of age. As expected, 24-month-old lower hindlimb muscles exhibited several characteristics of denervated muscle, including fragmentation of postsynaptic endplates, thinning of pre-synaptic motor axons, and increased varicosities in presynaptic endplates (**Fig. 7A, S10**). Subsequently, we applied the same 7-day SNC injury to aged (20-24 months) mice and performed similar myogenesis assays as young animals. Unlike young MuSCs that show increased cell volume, a characteristic of G_alert_ state, denervated MuSCs did not exhibit a difference in size (volume) compared to control aged MuSCs (**Fig. 7B)**. Similar *to mdx* muscle, no changes in the number of MuSCs were observed after nerve injury (**Fig. 7C).** Furthermore, myogenic differentiation of denervated and control aged MuSCs displayed no statistically significant differences (**Fig. 7D)**, and modulations of mitochondrial dynamics and metabolic function were also unchanged in aged MuSCs after denervation when compared to controls (**Fig. 7E, F)**. Contrary to young muscle in which denervation injury enhanced mitochondrial ETC content, we did not detect changes in mitochondrial ETC protein level as measured by Western blot (**Fig. S11)**. To further verify whether nerve-influenced muscle regeneration was also altered in the aged muscle, we induced the same nerve and muscle composite injury as in young and *mdx* mice. As predicted, unlike young muscle, SNC injury plus cryo-injury showed a similar rate of regeneration as the cryo-injury only group (**Fig. 7G)**. Lastly, in stark contrast to young muscle, when the same donor GFP^+^ MuSCs were transplanted, the engraftment efficiency of aged muscle (24 months) was lower in the TA muscles that were injured concurrently (**Fig. 7H)**. Taken together, results from dystrophic and aged muscle indicate that healthy neuromuscular junctions, as observed in youth, are indispensable for synergistic enrichment in myogenic function and reinforces the importance of neural niche in MuSC dynamics.

**Figure 7.**
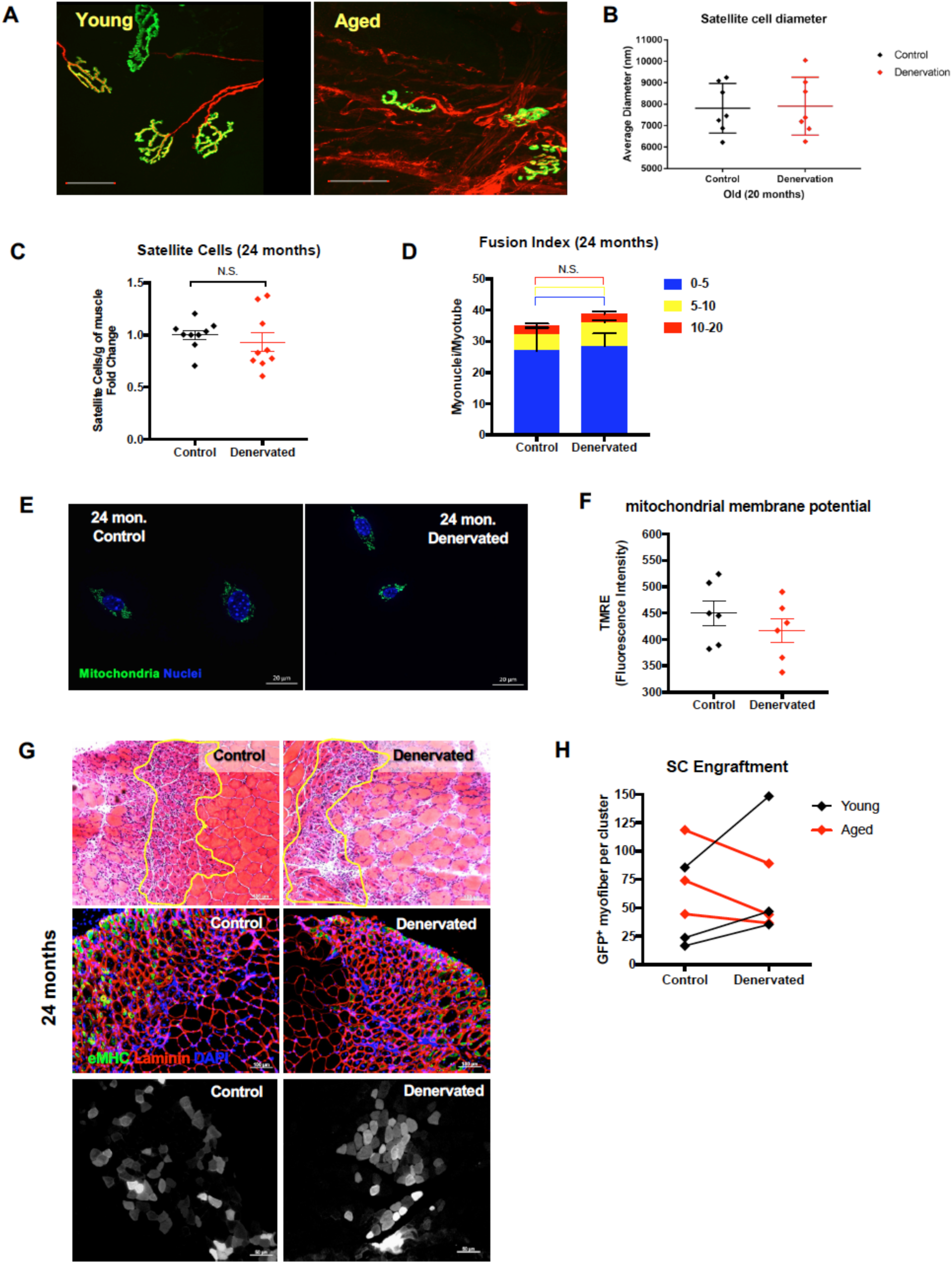
Satellite cell and motor neuron interactions are lost in aged muscle. **(A)** Representative images of neuromuscular junction from young (4 months) and old (24 months) extensor digitorum longus muscle. Presynaptic motor neuron is pseudo colored in red and postsynaptic acetylcholine receptors are depicted in green. **(B)** Quantification of FACS purified satellite cell diameters from 24-month-old control and denervated muscle. **(C)** Muscle satellite cell frequency from aged control and aged muscle 7 days following denervation. Data expressed as Mean ± SEM. **(D)** Representative images of aged control and denervated MuSC mitochondrial morphology during proliferation. **(E)** Quantification of fusion index (number of nuclei per myotube) from aged control and aged denervated MuSCs. n=4 **(F)** Mitochondrial membrane potential measured by TMRE in MuSC from aged control and aged denervation. **(G)** H&E (*top*), immunofluorescence staining (embryonic myosin heavy chain, laminin, DAPI (*middle*)), and satellite cell transplantation (*bottom*) cross section from 24-month-old TA muscle of 7-day cardiotoxin injury (right leg) and cardiotoxin + sciatic nerve pinch injury (left leg). **(H)** Quantification of transplantation efficiency comparing young control *vs.* denervated (black) and aged control vs. denervated (red). n=3 contralateral leg from same animal was used as control. **(I)** Representative Western blot image of mitochondrial electron transport chain subunits from lysates of control and denervated TA muscle. (Scale bars; (A) 50 µm, (E) 20 µm, (G) top, middle 100 µm, bottom 50 µm)

## DISCUSSION

The health and maintenance of skeletal muscle are mediated through functional MuSCs and the innervation by the neural network through NMJ, but interactions between MuSCs and the NMJ is not fully understood. In this study, we investigated the effect of NMJ perturbation on MuSC dynamics and observed healthy nerve regeneration transitions MuSCs into a “G_alert_” state, and boosts myogenic progression, possibly to rebuild muscle and compensate for the loss of mass. This enrichment in myogenesis is mediated by increased mitochondrial metabolism and upregulation of protein synthesis through activation of mTORC1. We also provide convincing evidence that synergistic interactions between the neuromuscular niche and MuSCs are abolished in pathological denervated conditions, such as advanced aging and Duchenne muscular dystrophy.

The reversible transition between quiescence and MuSC activation is critical for sensing perturbations to skeletal muscle homeostasis, but permanent insults such as denervation or from disease interrupt this balance and force eventual depletion of the functional MuSC pool (Dedkov et al., 2003) and atrophy (Dumont et al., 2015; Kuang et al., 2007). Previous work has shown that upon denervation, an increase in expression of synaptic and atrophic genes are induced in skeletal muscle myofibers (Luo et al., 2019), and this program is, in part, mediated through upregulation of Myogenin and Histone deacetylase 4 (HDAC4) (Tang and Goldman, 2006). Our results here suggest the upregulation of a proteolytic program in transiently denervated myofibers are offset by a retrograde protein synthesis program in MuSCs, underscoring the importance of MuSCs in functional recovery following denervation. We postulate that in long-term denervation injury, MuSCs engraft (Liu et al., 2015), to counteract the protein breakdown program from existing myonuclei. However, because the number of newly MuSC-derived myonuclei is much lower than existing myonuclei, a loss of muscle mass and pathological remodeling ensues. Additionally, crosstalk between existing and new MuSC-derived nuclei in the syncytium may further attenuate the protein synthesis program. Integrating these results suggest MuSCs engraft without myofiber damage in response to synaptic perturbations as a mechanism to resist pathological remodeling/adaptation.

We speculate several sources that signal to MuSCs to transition into a “G_alert_” state, which may act independently or concurrently in response to nerve regeneration, and be diminished by aging or disease. Schwann cells are potent sources of neurotrophic factors, and in aging, responses of Schwann cells after peripheral nerve injury are diminished (Painter et al., 2014). Further understanding of Schwann cell – MuSC crosstalk is warranted, but in response to nerve injury, Schwann cells emit a series of neurotrophic factors that have been shown to exert potent effects on MuSCs. For example, brain-derived neurotrophic factor (BDNF) and its receptor, p75^NTR^ are expressed in Pax7^+^ MuSCs and are known to regulate myogenic differentiation and muscle regeneration (Clow and Jasmin, 2010; Miura et al., 2012). In addition to its role as a trophic factor in motor neurons, BDNF has been identified as a key exercise-induced myokine that improves energy homeostasis (Delezie and Handschin, 2018; Koliatsos et al., 1993). Glial-derived neurotrophic factor (GDNF) and its subfamily of ligands, neurturin (NRTN), artemin (ARTN), and persephin (PSPN), which bind to the glycosylphosphatidylinositol-anchored GFRα receptors 1 through 4, are also neuroprotective and regenerative factors (Barati et al., 2006; Shvartsman et al., 2014; Walker and Xu, 2018; Zhang et al., 2009). Recently, supplementation of GDNF (a Ret ligand) was observed to rescue age-associated defects in MuSCs (Li et al., 2019), further supporting the positive role of neurotrophic factors, such as GDNF, in myogenesis. Consistent with these observations, increased Pax7^+^ MuSC content from young 7-day denervated muscle was accompanied by a 4-fold increase in GDNF mRNA, upregulation of nerve growth factor receptor (NGFr), which has been shown to demarcate myogenic progenitors with enhanced regenerative potential (Hicks et al., 2018), and GDNF family receptors (GFRα1 and GFRα3) (**Fig. S7**). These results suggest increased interactions between nerve trophic factors and MuSC state are critical in reestablishing and remodeling the neuromuscular synapse.

The perturbation of the neuromuscular synapse has distinct consequences for shaping the local muscle microenvironment through both activations of new cellular programs and metabolic rewiring. Cell types that respond to denervation such as FAPs have been shown to emit interleukin 6 (Madaro et al., 2018), a pleiotropic cytokine that is associated with atrophy (Munoz-Canoves and Michele, 2013), and in aging, secrete interleukin 33, an alarmin associated with environmental stress, as well as display preferential interactions with nerve structures such as muscle spindles (Kuswanto et al., 2016). Furthermore, it has been well documented that immune cells, especially myeloid cells, play a pivotal role in peripheral nerve regeneration (Barrette et al., 2008; Cattin et al., 2015; Chen et al., 2015; Mueller et al., 2003; Stratton et al., 2018). Upon nerve injury, macrophages accumulate at the site of the lesion, release vascular endothelial growth factor A (VEGF-A) and other cytokines that regulate maturation of Schwann cells, and are educated by the local microenvironment to polarize towards an anti-inflammatory phenotype (M2). The mitogenic activity of different growth factors and cytokines from activated FAPs and macrophages on MuSCs has been well established, and both have been shown to stimulate MuSCs to enter the cell cycle and indelibly change the availability of metabolites in tissue. However, in aging, the cellular and molecular complexion of these cells are altered and contribute to variations in MuSC crosstalk (Lukjanenko et al., 2019).

The anatomical positioning of MuSCs between the basal lamina and sarcolemma makes this population of stem cells particularly sensitive to calcium signaling. Given its indispensable role in synaptic transmission and muscle contraction, Ca^2+^ activity could be the key initial trigger of cellular communication following nerve injury. In support of this notion, a recent study revealed that denervation-induced alterations in Ca^2+^ signaling in muscle signals HDAC4 to shuttle between the nucleus and cytoplasm. HDAC4 has been shown to promote MuSC proliferation (Marroncelli et al., 2018) and co-localize with sub-synaptic myonuclei after denervation (Castets et al., 2019). Whether similar Ca^2+^-mediated mechanisms of myogenesis modulate MuSCs after denervation is unclear, but the calmodulin (CaM)-dependent kinase (CaMK), which prevents the formation of HDAC-myocyte enhancer factor 2 (HDAC-MEF2) complexes, releases MEF2 from HDAC4 and HDAC5 and promotes myogenic transcriptional activity in response to Ca^2+^ activity. Based on our transcriptome data, we hypothesize that nerve injury enriches MuSCs for hypertrophic growth as a compensatory mechanism to counterbalance atrophic signal caused by disruptions in Ca^2+^ dynamics in denervated myofibers. Furthermore, given the well-established sarcolemmal membrane damage and disruption in Ca^2+^ homeostasis manifested in DMD, as well as in aging, our results point to Ca^2+^ dynamics and HDAC4 as a potential therapeutic target to restore nerve-MuSC interactions in pathological denervation, such as in DMD and aging. Another important aspect of Ca^2+^ signaling and its potential link in regulating MuSC function is Ca^2+^-induced modulation of mitochondrial metabolism. Mitochondria function as secondary storage of Ca^2+^ to endoplasmic/sarcoplasmic reticulum and control various Ca^2+^ dependent signaling. Uptake of Ca^2+^ by mitochondrial Ca^2+^ uniporter into intermembrane space and matrix activates several mitochondrial Kreb cycle enzymes to promote oxidative phosphorylation and myogenic progression of MuSCs. Following SNC injury, we observed a significant upregulation in nuclear-encoded mitochondrial genes, and this upregulation was coupled with enhanced mitochondrial bioenergetics. Given perturbations in Ca^2+^ uptake in mitochondria are directly associated with the production of reactive oxygen species (ROS) and oxidative damage, which inhibit MuSC functions, these data suggest disruptions to Ca^2+^ signaling may act as a relay within the neuromuscular niche to stimulate MuSC function as a compensatory mechanism.

Our results provide strong evidence of synergistic interactions between neural circuitry and MuSCs, and that chronic disruption or degeneration of neuromuscular synapses, such as in muscular dystrophy and biological aging, significantly diminishes this synergy. These results underscore the importance of neuromuscular junction and neural network as an essential niche of muscle stem cells and provide a viable target for interventions against severe muscle trauma (Aguilar et al., 2018; Anderson et al., 2019; Mohiuddin et al., 2019), sarcopenia, and neuromuscular disorders.

## EXPERIMENTAL PROCEDURES

### Mice

All animal procedures were conducted under the approval of the Institutional Animal Care and Use Committee (IACUC) of the Georgia Institute of Technology and performed in accordance with all relevant guidelines and regulations. All mice in this study were either C57BL/6J genetic background or backcrossed with C57BL/6J for more than 6 generations and were initially purchased from Jackson Laboratory. Mice were bred and maintained in pathogen-free conditions with a 12-12 light/dark cycle in the Physiological Research Laboratory (PRL) at Georgia Institute of Technology. For muscle satellite cell reporter, mice expressing a tamoxifen-inducible Cre from the endogenous Pax7 promoter were bred with mice carrying a loxP-flanked STOP cassette followed by *TdTomato* or enhanced yellow fluorescent protein (EYFP) in the ROSA26 locus. Alternatively, *Pax7-ZsGreen* mice were used for FACS purification. Other transgenic reporter mice include: *Thy1-EYFP* (line 16) for motor neuron(Feng et al., 2000), mitoDendra2 for mitochondria(Mishra et al., 2015; Pham et al., 2012), *mdx-4CV* mice were used for mouse model of Duchenne muscular dystrophy (DMD). Females aged between 3 and 6 months, considered young adults and 22 to 24 months were consider aged group. Mice were randomized for all experiments in this study. No differences in muscle regeneration were observed in response to denervation injury between 3-month old and 6-month old mice.

### Animal injury models

Mice were anesthetized by inhalation of 2.5% isoflurane. The bare skin above the sciatic nerve was disinfected with 70% ethanol and chlorhexidine swaps. The disinfection step was repeated 3 times with 70% ethanol and chlorhexidine followed by 70% ethanol scrubbing. A small opening of the skin was performed to expose a sciatic nerve. The exposed nerve is then crushed with forceps for 10 seconds. The contralateral leg was served as a sham control. Wound clips were used to close the open incision and they were removed 14 days after the surgery.

### Muscle stem cell isolation

MuSCs were isolated through a cell sorting procedure as previously performed (Cerletti et al., 2008; Han et al., 2018). Briefly, hindlimb muscle tissues were harvested from young (2-4 months of age) and aged (20-25 months of age) mice, and then they were incubated in 20 mL of DMEM media containing 0.2 % collagenase type II and 2.5 U/ mL dispase for 90 min at 37 °C. After tissue digestion, the resulting media were mixed with same volume of stop media (20% FBS in F10), filtered using a cell strainer with a pore size of 70 μm, and then centrifuged (300 g for 5 min. at 4 °C) (Allegra X-30R Centrifuge, Beckman Coulter, USA) to obtain the myofiber-associated cell pellet. The cells pellets were washed with HBSS containing 2% donor bovine serum (DBS), and the cells were incubated with primary antibodies. For MuSC sorting, a cocktail mixture containing the following antibodies was used: (1) allophycocyanin (APC)-conjugated anti-mouse CD11b (1:200), CD31 (1:200), CD45 (1:200), Sca-1 (1:200), and Ter119 (1:200), (2) phycoerthrin (PE)-conjugated anti-mouse CD29 (1:100), and (3) biotinylated anti-mouse CD184 (1:100). After incubation for 30 minutes at 4 °C, the primary antibodies-treated cells were washed, centrifuged (300 *g* for 5 minutes at 4 °C), and then treated with a secondary antibody (Streptavidin PE-Cy7) (1:50) for 20 minutes at 4°C. Following propidium iodide (PI) treatment and strainer filtration (70 μm), the MuSCs (negative selection: PI, CD11b, CD45, Sca-1, and Ter119; positive selection: CD29, β1-integrin and CD184, CXCR4) were isolated using fluorescence activated cell sorting (FACS) (BD FACS Aria III, BD Biosciences, USA). The freshly sorted MuSCs were used without sub-culturing.

### MuSC transplantation

Tibialis anterior (TA) muscles of recipient mice were pre-injured with BaCl_2_ 24 hours prior to the MuSC transplantation. On the day of transplantation, FACS-isolated MuSCs (15,000∼20,000 cells) were suspended in PBS and injected intramuscularly to the TA. A small incision of skin was made to expose the muscle. The injection was performed with a Hamilton syringe (VWR 72310-316) and the injection volume of the cell suspension was 10 μL. The needle was injected into the distal TA muscle. After the injection, the incision was closed by wound clips.

### Oxygen consumption rate

XF Cell Mito Stress tests were performed on FACS purified MuSCs. Briefly, 10,000 MuSCs per well were seeded on Seahorse miniplate. Sensor cartridges were incubated in XF Calibrant overnight in 37°C. The following day, the cells were placed in non-CO_2_ incubator at 37°C for an hour in assay media. The sensor cartridge was then placed in XFp Analyzer for calibration. When calibration was done, cartridge was removed and replaced with the cell miniplate for mito stress test. Basal oxygen consumption rate were measured and then the mitochondrial stress test was initiated by adding different mitochondrial inhibitors in ports, A, B, and C were injected with 10 µM oligomycin, 5 µM CCCP, and 5 µM rotenone/antimycin A, respectively. After the assay, the cells were lysed in RIPA to calculate protein concentration for further normalization.

### Histology and immunohistochemistry

Whole hindlimbs were collected and fixed with 4% paraformaldehyde for 3 hours. The fixed legs were washed 3 times for 5 minutes with PBS and then TA muscles were dissected. The muscles were incubated in 20% sucrose in PBS overnight at cold room. Following PBS washing, the tissues were mounted on cork with OCT compound (Electron Microscopy Sciences 62550-01) to be frozen for the cryosection. Tissues were cut to 10 μm thickness and placed on slide glasses. Before the antibody staining on the muscle section, blocking was done with blocking buffer (2% BSA, 0.5% Gt serum, 0.5% TritonX-100 in PBS) for 30 min at RT followed by another blocking using M.O.M kit (Vector Laboratories BMK-2202). Primary antibodies (mouse anti-MYH3; DSHB F1.652-s, rabbit anti-Laminin; Sigma L9393) were incubated overnight in a cold room followed by washing with PBS-T (0.1% Tween-20 in PBS) 3 times. Then, secondary antibodies (Goat anti-mouse IgG1 Alexa Fluor 488 conjugate; Invitrogen A21121, Goat anti-rabbit H+L Alexa Fluor 555 conjugate; Invitrogen A21428) with Hoechst 33342 (Thermo H3570) were incubated for 2 hours in room temperature. After washing the muscle with PBS-T twice and PBS once, a drop of mounting medium with DAPI (Abcam ab104139) was added on the top of the sample and covered with a cover glass. Antibodies were diluted to the suggested dilution ratio following the manufacturers’ instruction.

### Western blot analysis

Gastrocnemii were homogenized with 25 strokes using a 5 mL PTFE tissue grinder with clearance 0.15-0.25 mm (VWR 89026-392, 89026-404) at 3,000 rpm in RIPA lysis buffer (VWR 97063-270) supplemented with Roche cOmplete Mini Protease Inhibitors (Roche 04693124001) and PhosSTOP Phosphatase Inhibitors (Roche 04906837001). Following 3 freeze-thaw cycles in liquid nitrogen and on ice, the samples were centrifuged at 18,400 *g* for 10 minutes, and the supernatants (homogenates) were normalized to total protein concentration using a BCA protein assay kit (Thermo 23225). 50 µg of protein were run through 4-20% Criterion TGX Gels (Bio-Rad 5671093) at 150 V for 185 minutes and transferred to a PVDF membrane using a Trans-Blot Turbo System at 2.5 A for 7 minutes. Ponceau staining (Sigma P7170) was used as a loading control. Membranes were imaged on Li-Cor Odyssey CLx-1050 Infrared Imaging System and bands were quantified on Li-Cor Image Studio V5.2.

### RNA Isolation and Quantitative Polymerase Chain Reaction

Using the normalized tissue homogenate prepared for western blot analysis, 50 µL of homogenate was added to 300 µL of RLT buffer supplied in the RNeasy® Mini Kit (Qiagen 74104) supplemented with 1% β-mercaptoethanol to inactivate RNases. The protocol according to Qiagen RNeasy® kit was followed for the remaining RNA isolation steps. The RNA concentration was measured by a NanoDrop One while A260/230 and A260/280 ratios were calculated for quality control. RNA content was then calculated by normalizing the RNA concentration to total muscle mass. To reverse transcribe the RNA to copy DNA, RNA concentrations were normalized to each other and the protocol according to Applied Biosystems High-Capacity cDNA Reverse Transcription Kit (Applied Biosystems 4368814) was followed, and the samples were run in a thermal cycler according to the recommended conditions. Finally, the Applied Biosystems PowerUp SYBR Green Master Mix (Applied Biosystems A25742) was used with the primers found in Supplementary Table S4 in an Applied Biosystems StepOnePlus Real-Time PCR system to perform the qPCR reactions. β-actin and B2M, which were found to be stably expressed following ischemia, were used housekeeping genes to quantify relative fold induction.

### Transmission electron microscopy

We fixed muscle block (1 mm^3^ in size) with 2% paraformaldehyde and 2.5% glutaraldehyde in 0.1 M sodium cacodylate buffer, post-fixed with 1% osmium tetroxide followed by 1% uranyl acetate, dehydrated them through a graded series of ethanol washes and embedded them in resin. Blocks were cut in ultrathin (80 nm) sections on Reichert Ultracut UCT, stained the sections with uranyl acetate followed by lead citrate, and viewed on a JEOL 1230 EX transmission electron microscope at 80 kV.

### Statistical analysis

Quantitative data are expressed as mean ± SEM. Statistical analysis was performed using GraphPad Prism 8, and statistical comparisons of analyzed values were obtained from *t*-test, one-way or two-way analysis of variance (ANOVA). For multiple group comparisons, one-way ANOVA with Tukey’s post hoc tests or Dunnett’s post hoc tests and two-way ANOVA with Sidak’s post hoc tests were performed. All tests resulting in *p*-value (<0.05) were considered statistically significant.

### Accession codes

RNA-Seq data from this study has been archived at the NCBI Sequence Read Archive under BioProject [PRJNA:625722].

## ACKNOWLEDGEMENTS

We thank the Physiological Research Laboratory and the core facilities at the Parker H. Petit Institute for Bioengineering and Bioscience This work was supported by the National Institutes of Health under award numbers R21AR072287 (Y.C.J.), Department of Defense W81XWH-20-1-0336 (Y.C.J. and C.A.A.), and S&R Foundation (Y.C.J.). The content is solely the responsibility of the authors and does not necessarily represent the official views of the National Institutes of Health.

## AUTHOR CONTRIBUTIONS

J.J.C., C.A.A., A.J.W., and Y.C.J. designed the experiments, analyzed the data, and wrote the manuscript. J.J.C., E.S., M.M., N.H.L., H.J., A.S., S.E.A., W.M.H., T.T., S.N., G.J., D.O., L.D.W., S.B.S., M.E.O., L.K., T.N.R., and C.A.A. conducted the experiments, contributed to methodology, and analyzed the data. L.W., C.A.A. provided significant contributions to methodology and conceptual justification. All authors reviewed the manuscript.

## SUPPLEMENTAL INFORMATION

**Figure S1 (related to Figure 1).**
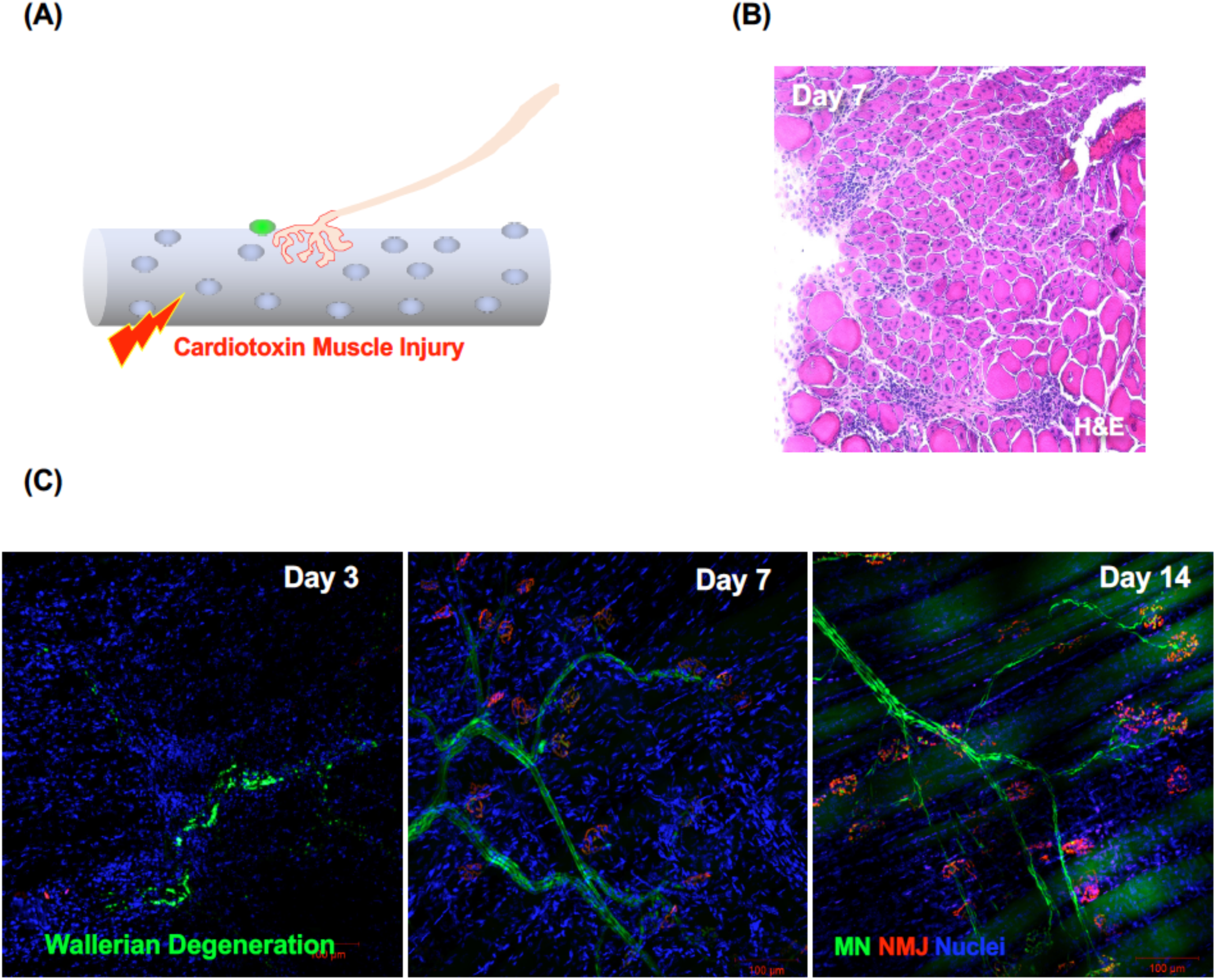
Motor unit is altered following cardiotoxin injury. **(A)**Schematic diagram depicting cardiotoxin injury. **(B)** Representative image of H&E staining 7 days after cardiotoxin injury. **(B)** Representative images of peripheral nerve regeneration following cardiotoxin injury.

**Figure S2 (related to Figure 2).**
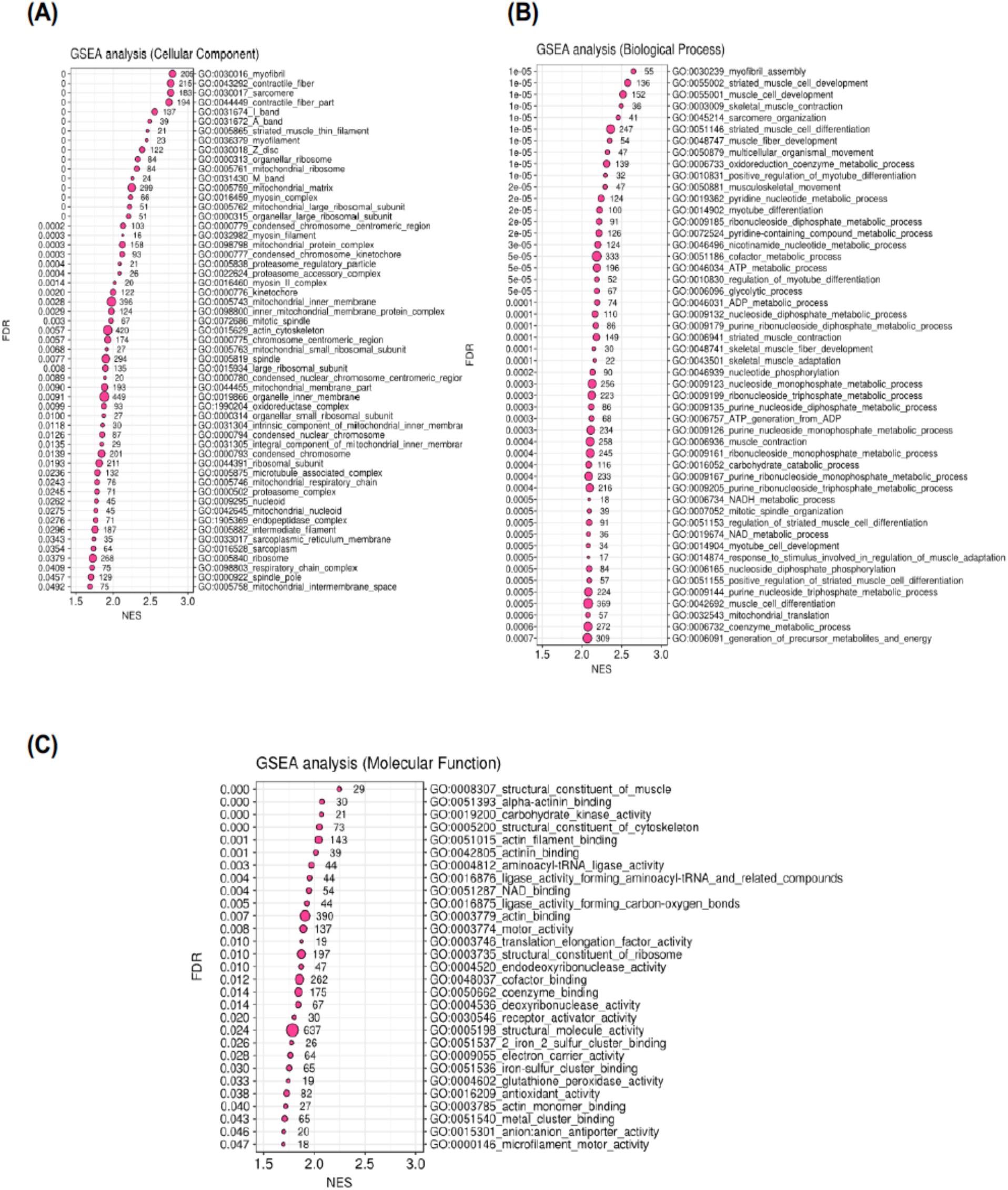
Transcriptomic analysis of muscle satellite cells following 7-day denervation. Gene set enrichment analysis (GSEA) of various pathways (cellular components, biological process, and molecular function shown) differentially regulated in denervated muscle satellite cells 7 days following denervation.

**Figure S3 (related to Figures 3):**
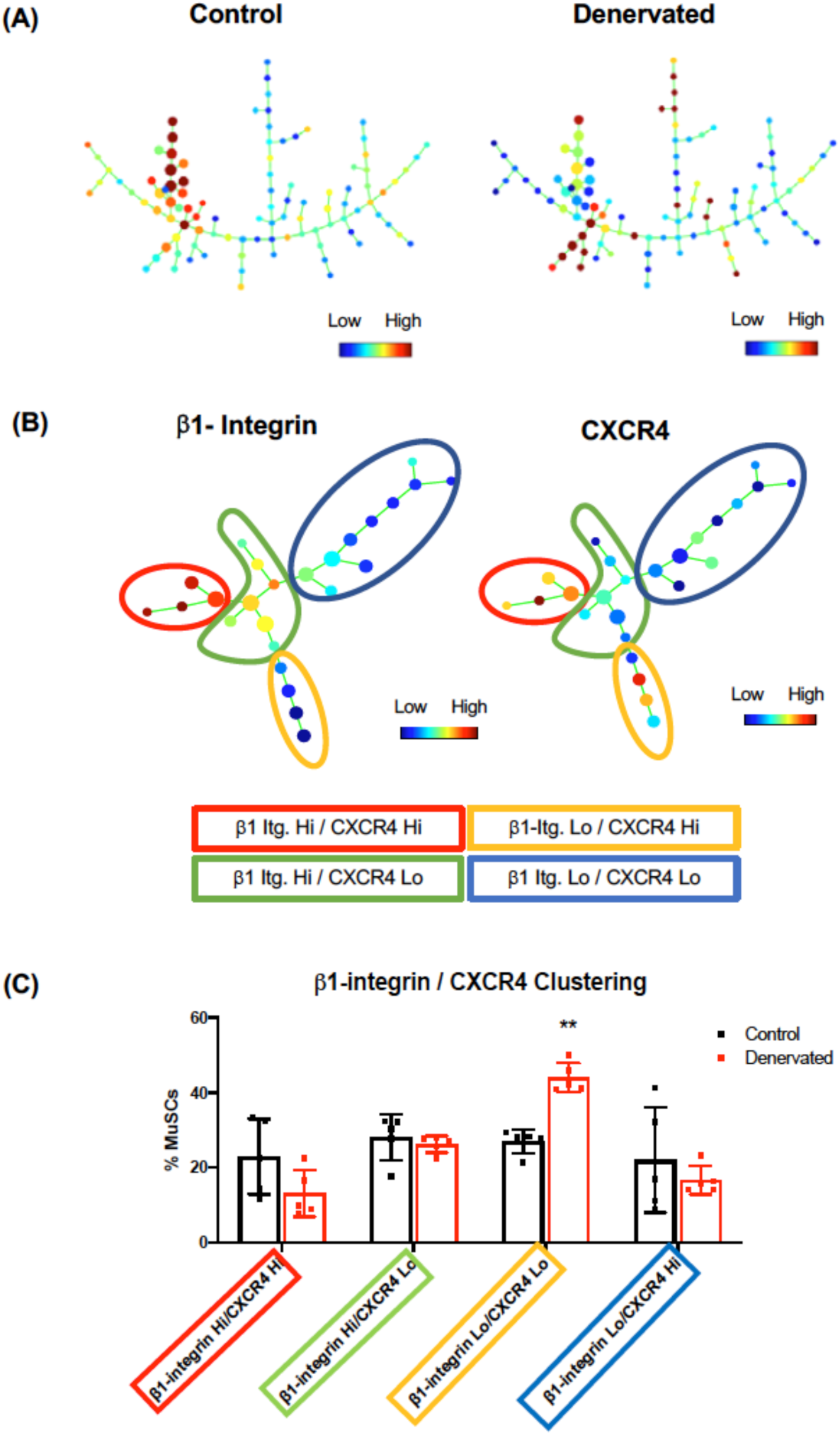
SPADE analysis of muscle satellite cells. **(A)** Representative SPADE tree demonstrating the frequency of cells from muscle tissue purified from control or 7 day denervated muscle. Total live cells were clustered on muscle stem cell markers (CD45^−^/CD31^−^/CD11b^−^/Ter119^−^/Sca1^−^/β1-integrin^+^/CXCR4^+^). **(B)** β1-integrin and CXCR4 marker expression levels are overlaid on SPADE diagrams of pre-gated MuSC showing variation in the expression of β1-integrin and CXCR4 among MuSCs. Images show representative results of five independent biological replicates. Nodes in both diagrams were grouped based on the distinct molecular features of each population. **(C)** Quantification of the node distribution of MuSCs from control or denervated muscles. Denervated muscle has significantly more β1-integrin Low/CXCR4 Low cells compared to control. \*\**p*<0.01, Mean ± SD

**Figure S4 (related to Figure 3).**
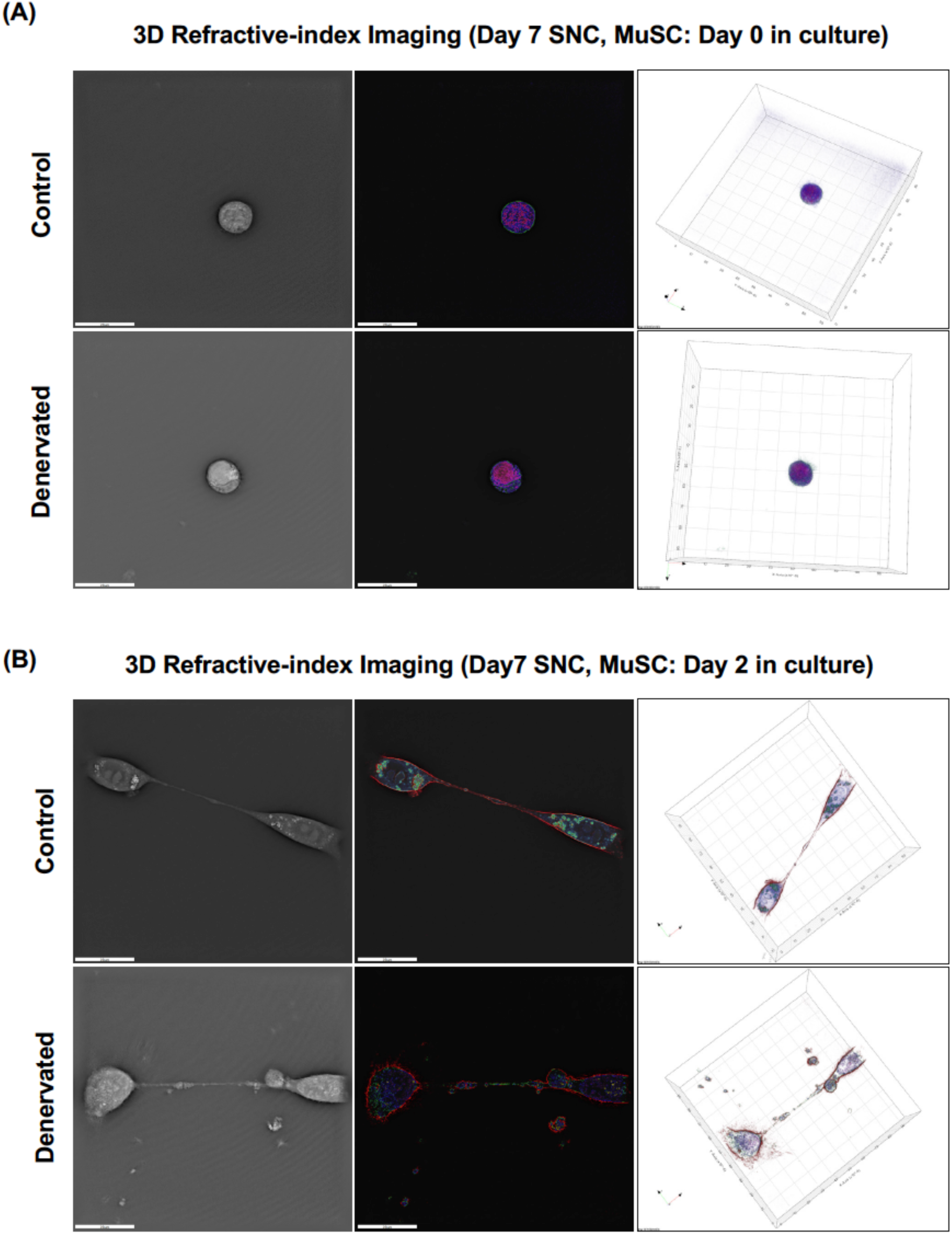
3D refractive-index imaging of muscle satellite cells. Representative capture of non-labeled, 3D holographic images of freshly isolated satellite cell depicting quiescent state **(A)** and myoblast fusion **(B)** from control and 7-day denervated muscle satellite cells. 3D refractive indices were pseudo-colored, Blue: nuclear components, Green: cytoplasmic components, and Red: membrane.

**Figure S5 (related to Figure 4).**
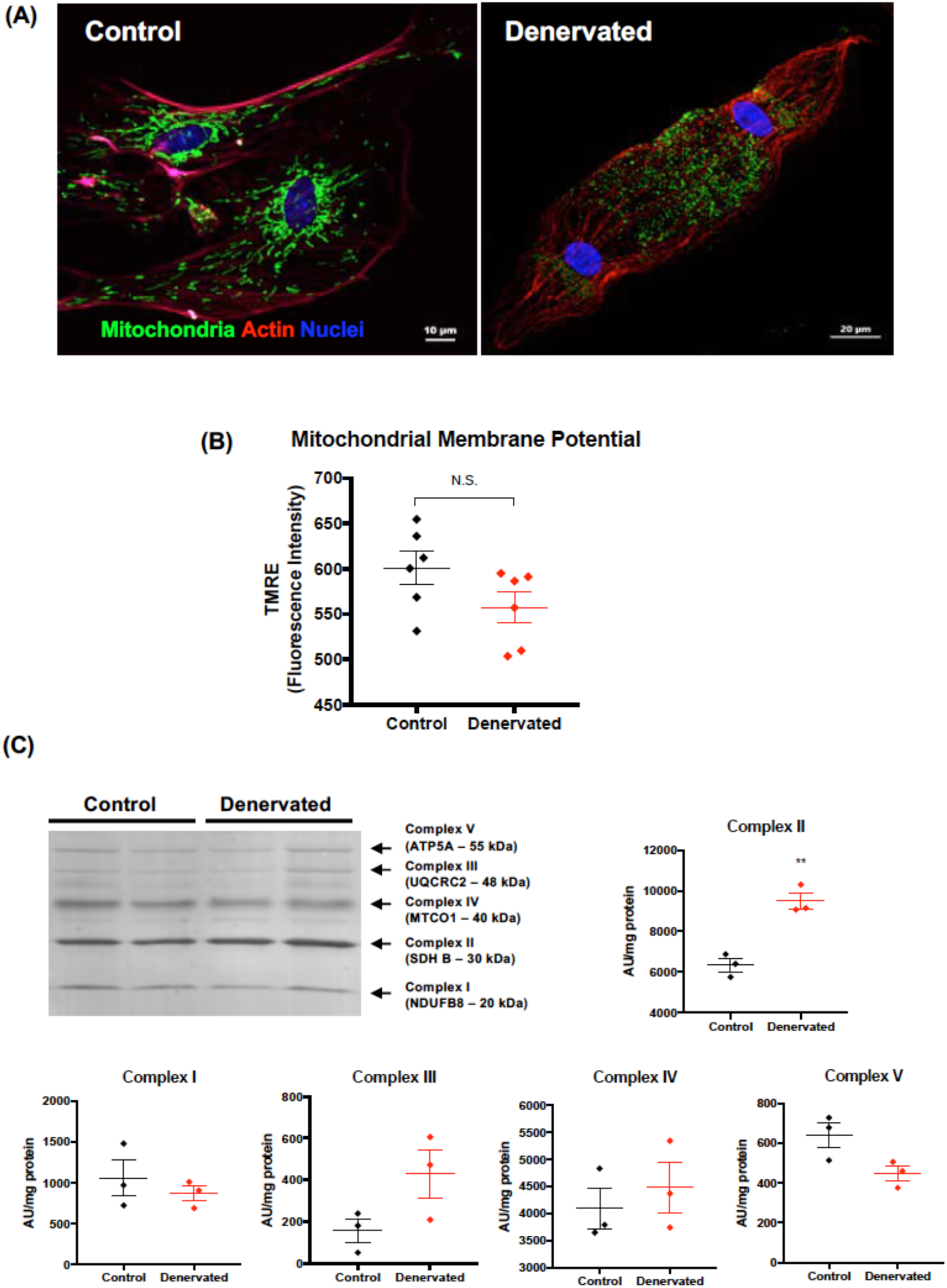
Mitochondrial functions are enhanced in denervated muscle satellite cells. **(A)** Representative images of muscle satellite cell mitochondria *ex vivo*. MuSC were isolated from transgenic mice that ubiquitously expresses Dendra2 fluorescence protein in mitochondrial matrix. Green: Dendra2, mitochondria, Red: F-actin, Phalloidin, and Blue: nucleus, DAPI. **(B)** Mitochondrial membrane potential measured by TMRE in MuSC from control and denervated satellite cells 5 days after seeding. (C) Representative Western blot image of mitochondrial electron transport chain subunits from lysates of control and denervated TA muscle. **(D)** Quantifications of mitochondrial electron transport chain subunit from each complex shown in **(C)**.

**Figure S6 (related to Figure 5).**
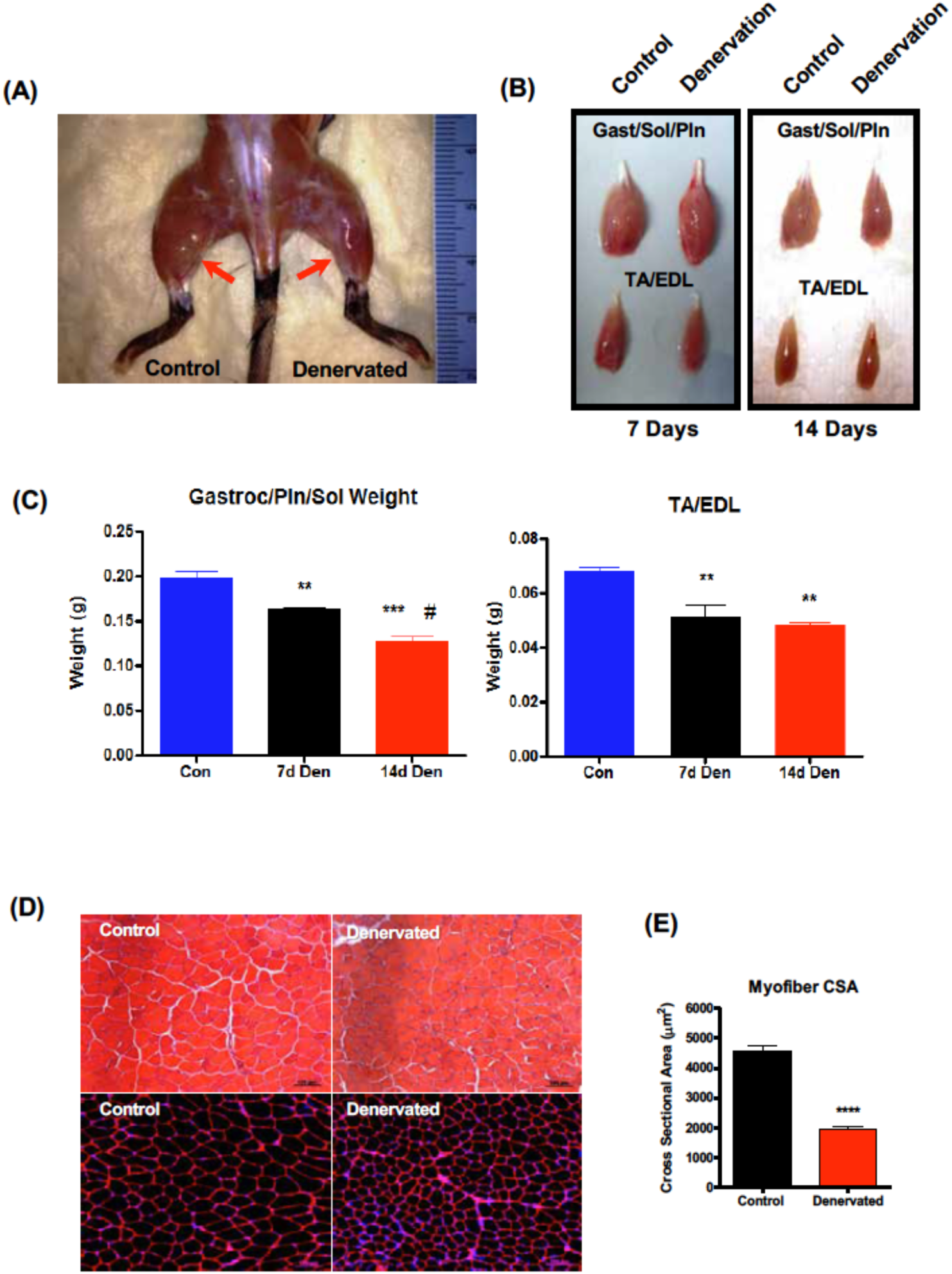
Morphological changes of skeletal muscle following sciatic nerve pinch injury. **(A)** Representative images of skeletal muscle exhibiting muscle atrophy. Red arrows indicate gastrocnemius muscle size difference between sham (control) and denervated 14 days after injury. **(B)** Comparison of lower hindlimb muscles, gastrocnemius (gastroc.), plantaris (Pln.), soleus (sol.), tibialis anterior (TA), and extensor digitorum longus (EDL) following 7 days and 14 days denervation injury. **(C)** Quantification of wet weight from lower hindlimb 7 days and 14 days after denervation injury. 14 days sham contralateral leg served as control. \*\**p*<0.01 \*\*\**p*<0.001, Mean ± SEM, n=4 **(D)** H&E (*top*) and immunofluorescence staining (laminin and DAPI) (*bottom*) cross section from TA muscle of 7-day after denervation. **(E)** Quantification of mean cross-sectional area. \*\*\*\**p*<0.0001, Mean ± SEM, n=4.

**Figure S7 (related to Figure 5).**
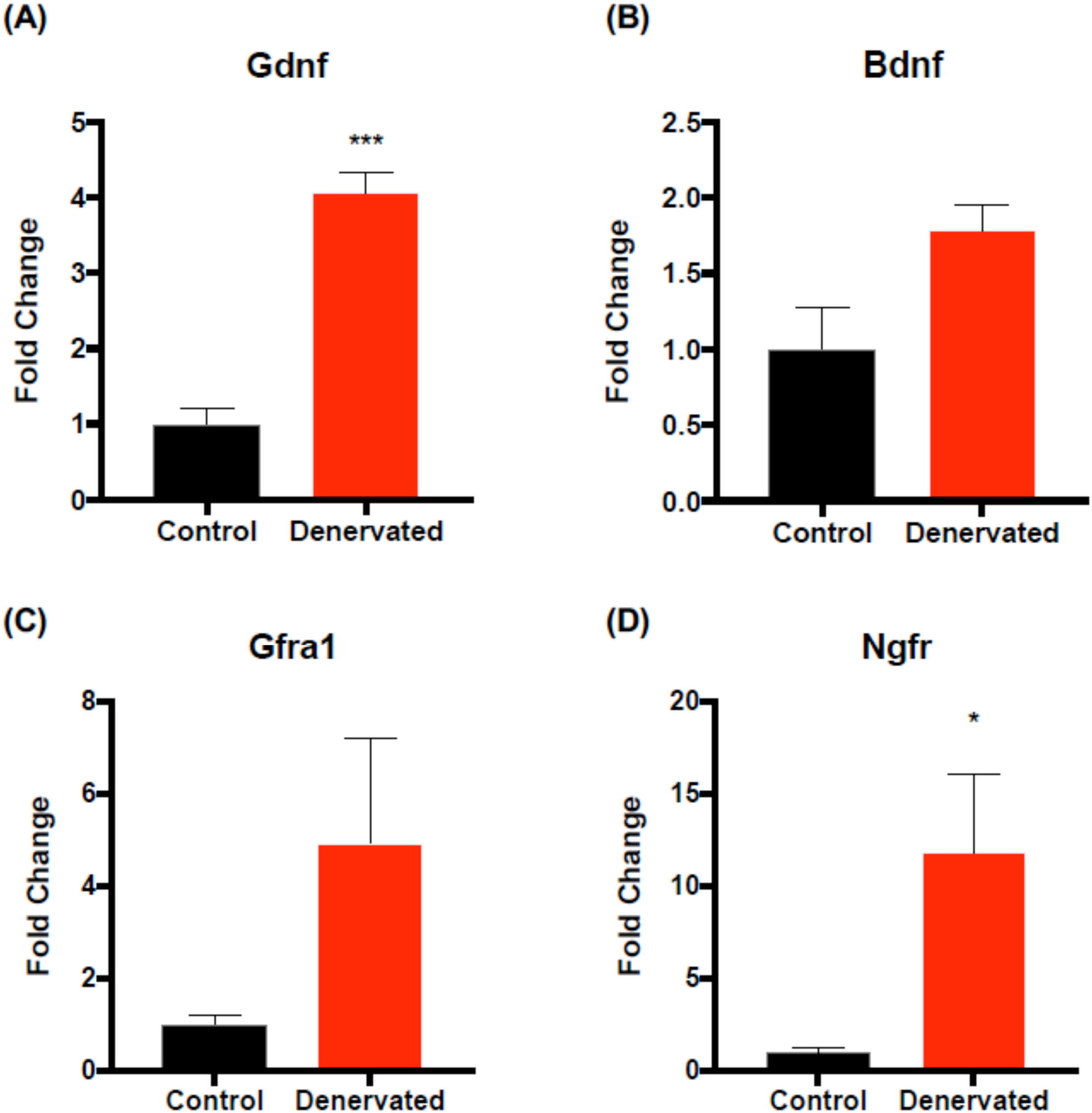
Neurotrophic factors are upregulated in denervated muscle satellite cells. Quantitative RT-PCR analyses of **(A)** glial-derived neurotrophic factor (Gdnf), **(B)** brain-derived neurotrophic factor (Bdnf), **(C)** GDNF family receptor a1 (Gfra1), and **(D)** nerve growth factor receptor (Ngfr) genes following 7 days denervation. \**p*<0.05, \*\*\**p*<0.001, Mean ± SEM, n=3

**Figure S8 (related to Figure 6).**
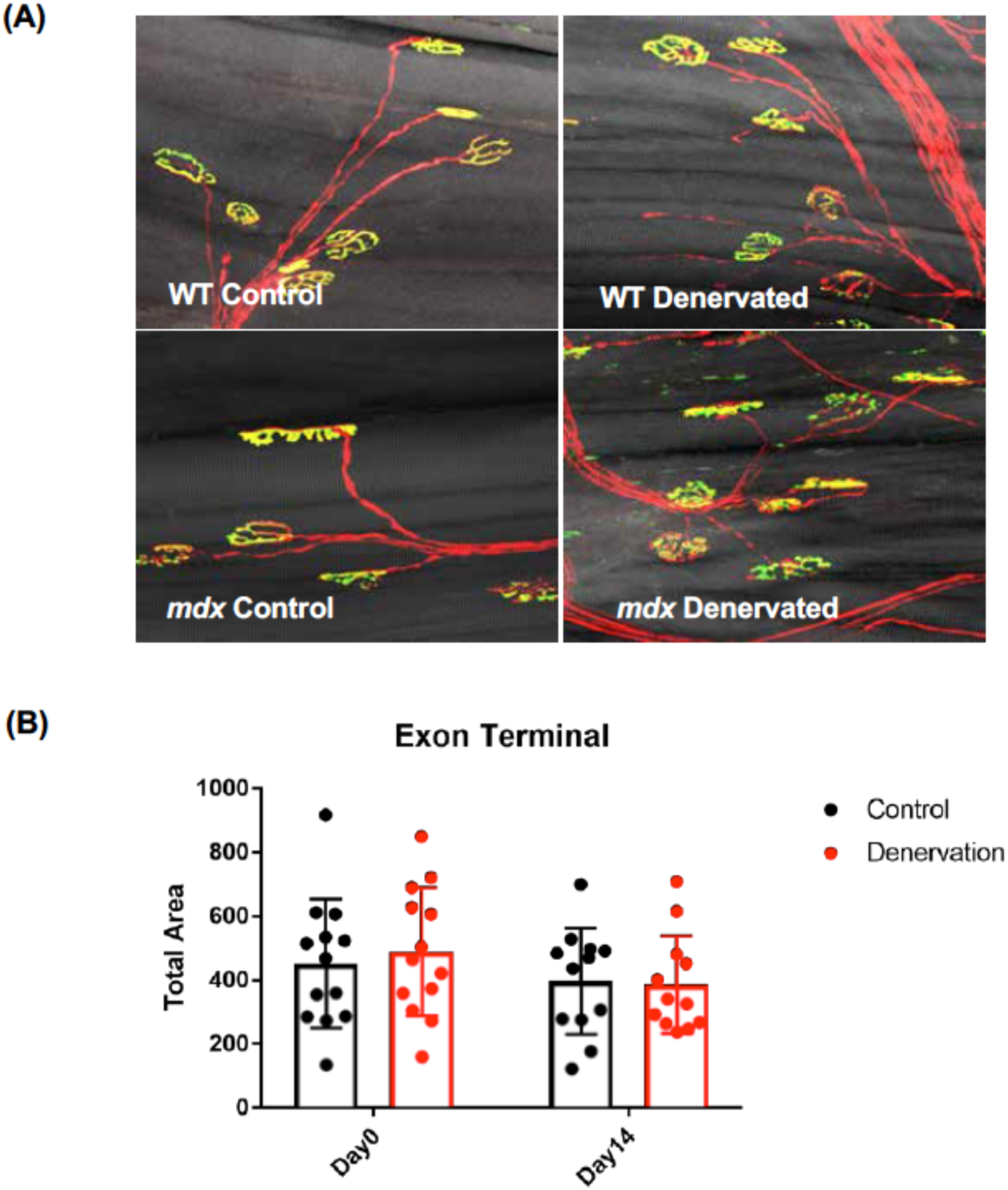
Peripheral nerve regeneration of *mdx* muscle. **(A)** Representative images of neuromuscular junction from wildtype (WT) and *mdx* old extensor digitorum longus muscle 14 days following sciatic nerve pinch injury. Presynaptic motor neuron is pseudo colored in red and postsynaptic acetylcholine receptors are depicted in green. **(B)** Quantification of neuromuscular junction area 0 and 14 days after injury.

**Figure S9 (related to Figure 6).**
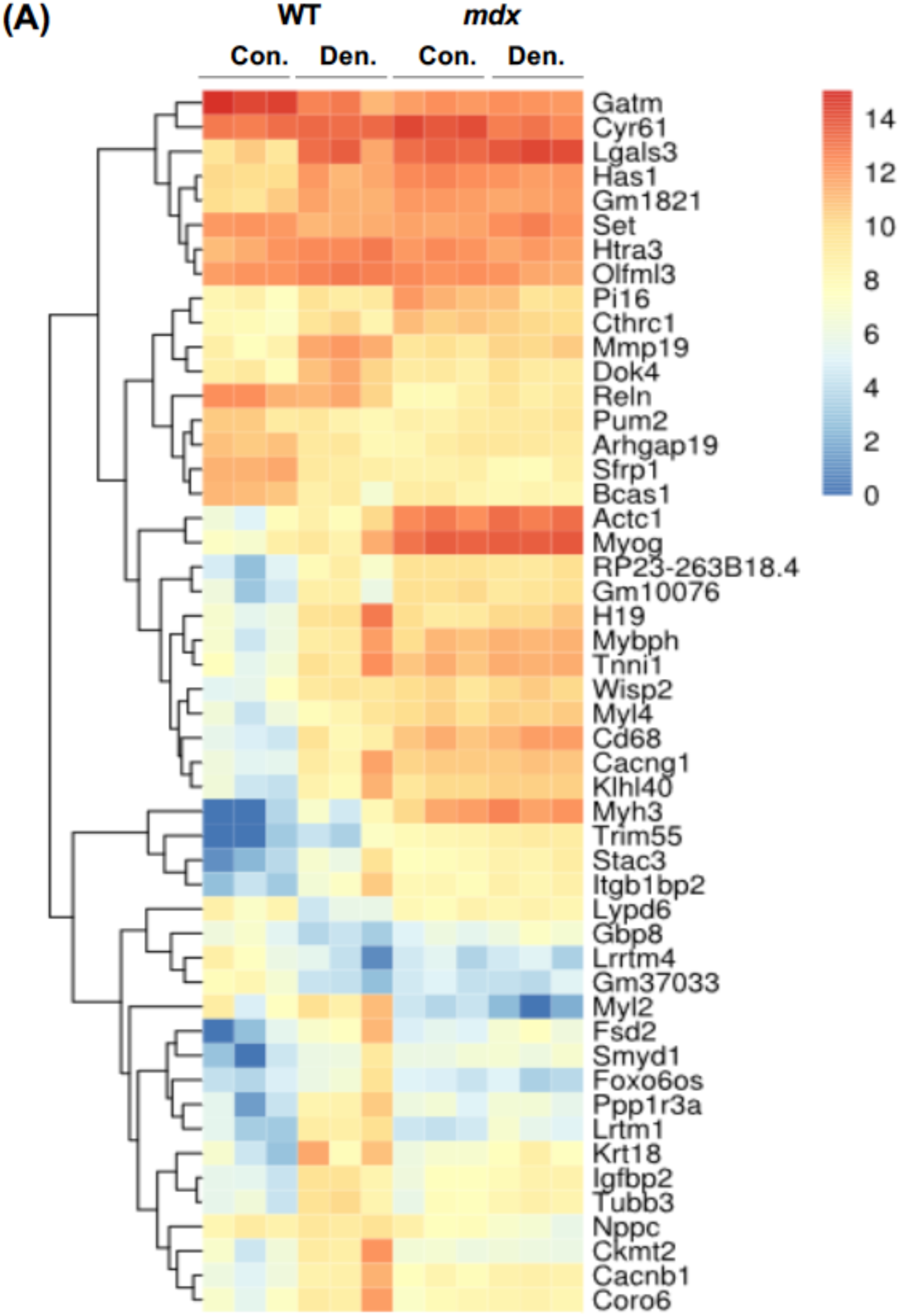
Peripheral nerve regeneration of *mdx* muscle. **(A)** Heatmap of top fifty differentially expressed gene clusters from WT and *mdx* following 7-day denervation.

**Figure S10 (related to Figure 7).**
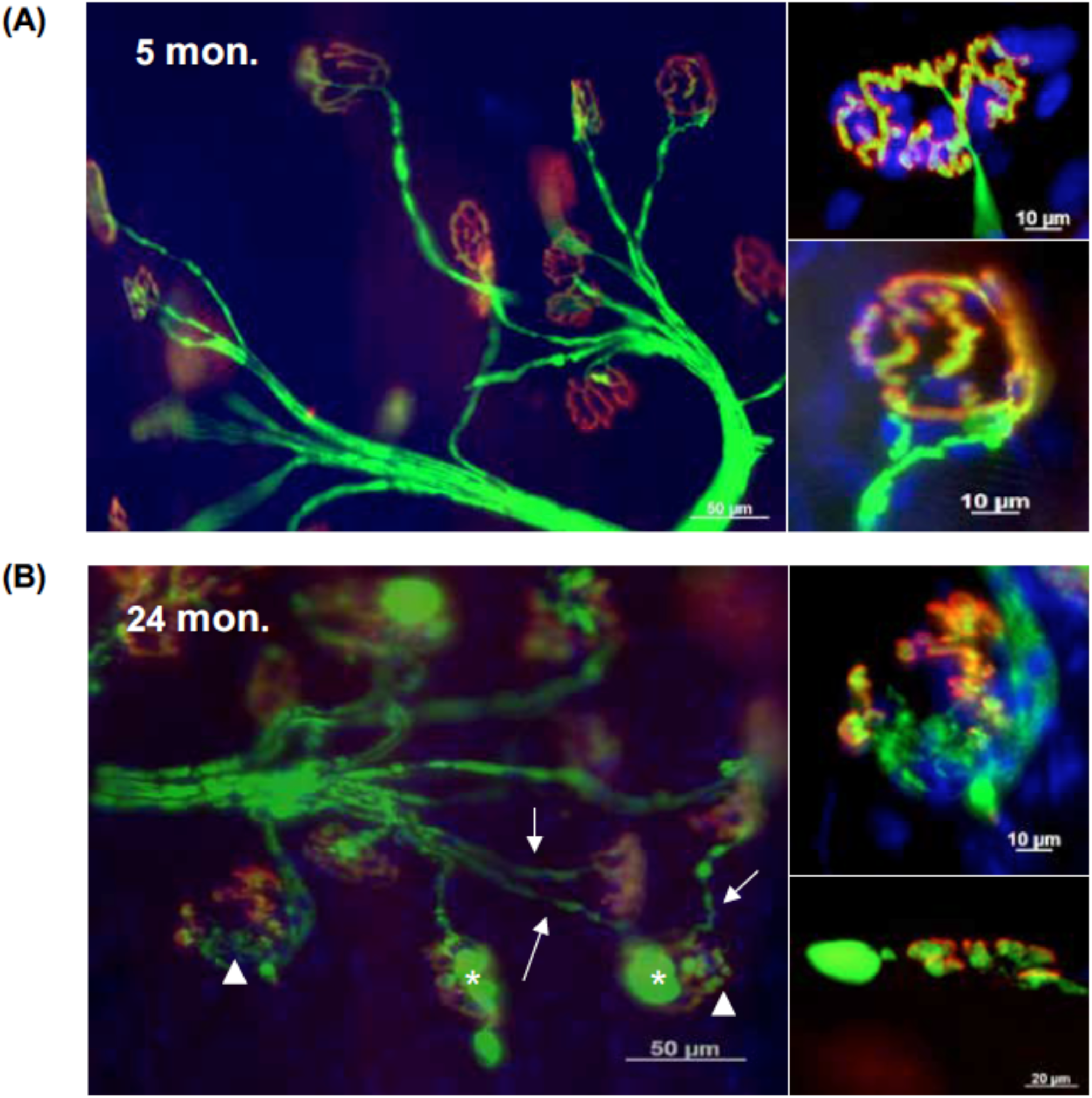
Neuromuscular junctions are degenerated in aged muscle. Representative images of neuromuscular junction from young (5 months) **(A)** and old (24 months) **(B)** extensor digitorum longus muscle. Presynaptic motor neuron is pseudo colored in green and postsynaptic acetylcholine receptors are depicted in red. Aged neuromuscular junctions exhibit a significant increase in fragmentation (arrow heads), thinning of presynaptic motor neuron (arrows), and bulged varicosities (*).

**Figure S11 (related to Figure 7).**
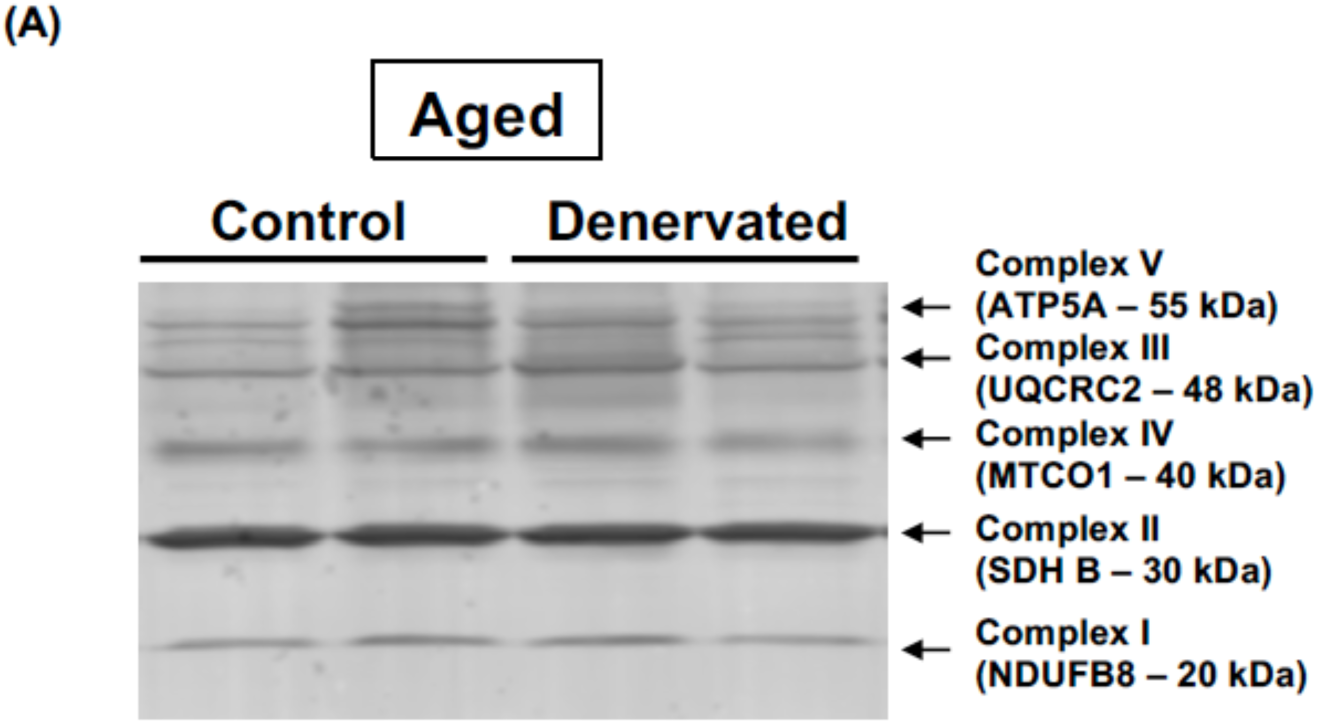
No change in mitochondrial electron transport chain content in aged muscle following nerve injury. **(A)** Representative immunoblot of mitochondrial ETC content

## Supplemental Methods

### mRNA Sequencing Library Preparation

mRNA was isolated and prepared into sequencing libraries as previously described^1^. Briefly, total RNA was isolated from sorted cells using Trizol (Thermo) and the miRNeasy Micro Kit (Qiagen) as per the manufacturer’s instructions and RNA concentration and integrity were measured with a Nanodrop spectrophotometer (Nanodrop 2000c) and Bioanalyzer (Agilent 2100). Approximately 10ng of isolated total RNA was used to produce cDNA libraries using the SmartSeq4 protocol (Clontech), as per the manufacturer’s instructions. Individual libraries were pooled and sequenced using 76-bp paired-end reads on an Illumina NextSeq to an average depth of 50M reads per library.

### Differential gene expression

Sequenced files were demultiplexed and aligned to the mm10 reference genome with the STAR algorithm (version 020201)^2^. Gene quantification was performed with RSEM (version RSEM-1.2.30)^3^ and replicates with Pearson correlation below 0.95 and genes with FPKM < 1 across all samples were excluded from analysis^4^.

The Bioconductor svaseq package (v 3.25.1) was used to perform surrogate variable analysis on the expected count field in the RSEM output (as proxy for raw read count). svaseq analysis was performed using a null model with explanatory variables of mouse phenotype (control vs injured). The lmfit function in R was used to perform a linear regression of each surrogate variable on the two explanatory variables – all surrogate variables were found to have *p*-value > 0.05 for the regression model fit. Principal component analysis was then performed on the R-log-transform of read counts in the individual biological replicates, using the R-log Transform function within the Deseq2 R package^5^. Surrogate variables were included as covariates in the linear model, resulting in a linear model with gene expression as output and input variables as control vs. injured, and 3 surrogate variables.

Differential gene expression analysis was performed with the bioconductor limma package (v 3.5)^5,6^. The voom function was used to obtain log2 counts per million and a linear model was then fit to the transformed counts. The eBayes function from the limma package was used to obtain fold changes and F-statistic values for the comparison. Genes were determined to be differentially expressed if the limma voom analysis determined a fold change greater than 2 with false discovery rate (FDR) corrected p-value below 0.01. The top Table function within limma voom was used to examine differential genes for injured *vs.* control.

### Differential pathways and GO terms

The DAVID software^7^ was used to identify significantly enriched KEGG pathways and GO terms for each set of differential genes. Differential gene lists were provided to DAVID to perform a functional clustering analysis with FDR cutoff of 0.01, and all other parameters were used at default settings. Additionally, differential gene lists for each of the four contrasts were loaded into Cytoscape (v.3.2.1)^8^, and the Reactome FI gene set/ mutation analysis app was used to generate a network of over-represented differential Reactome pathways (FDR threshold = 0.01). The differential genes were clustered with the EAGLE algorithm to identify subsets of genes that interact with one another.

### Analysis of differential long noncoding RNAs and transcription factors

Long noncoding RNA (lncRNA) differential expression analysis was performed using a list of known GRCm38 lncRNAs obtained from Entrez. The FPKM field of the RSEM gene expression profiles for the lncRNA’s with FPKM >=1 for at least one condition was analyzed. Replicates were averaged to get a mean expression value for each lncRNA for each comparison. Log2-fold change in expression was computed for injured vs control and abs (log_2_-fold change) >1 was used to determine whether a particular lncRNA was differentially expressed.

A similar approach was used to identify differentially expressed transcription factors. A list of known GRCm38 transcription factors was downloaded from Animal Transcription Factor Database^9^. Log2-fold change in FPKM was calculated for injured vs. control and abs(log2-fold) change cutoff of 1 was used to determine significance.

### Data visualization

Several approaches were used to visualize the data. The Washu Genome Browser^10^ was used to generate tracks of normalized RNA-seq counts for all conditions. The venn function in limma was used to generate a Venn diagram illustrating the intersection of the differential gene sets in each of these comparisons. The g-plots heatmap.2 function^11^ was used to generate row-scaled (Z-scaled) heatmaps of expression values for differentially expressed genes. Gene expression values were clustered using Euclidean distance across FPKM values in both the x-axis (condition of interest) and y-axis (FPKM value).

